# Arbuscular mycorrhizal symbiosis increases drought resistance in the xerophytic argan tree (*Sideroxylon spinosum*)

**DOI:** 10.64898/2026.06.20.733516

**Authors:** Abdellatif Essahibi, Laurent Falquet, Aurélie Esseiva, Ahmed Qaddoury, Ivan Mateus, Didier Reinhardt

## Abstract

The xerophyte argan (*Sideroxylon spinosum*) has great ecological and socioeconomic importance for Morocco. However, it is endangered due to climate change and human overexploitation. We assess drought resistance traits of argan and test the potential of arbuscular mycorrhizal (AM) symbiosis to promote its growth and mitigate the consequences of drought. We compare ten endemic Moroccan mycorrhizal inocula with the model AM fungus *Rhizophagus irregularis* and with the drought-adapted isolate *Diversispora omaniana*. We integrated physiological phenotyping and RNA sequencing to investigate the stress resistance mechanisms of argan against drought. We show that AM symbiosis significantly mitigates drought effects on plant growth, mainly by improving water relations and photosynthetic efficiency, resulting in increased growth rates. Taken together, physiological and transcriptomic analyses show that stress markers were moderatly induced during severe drought stress irrespective of mycorrhizal status, suggesting that argan adopts a drought-coping strategy that involves both, stress avoidance and stress tolerance. Argan is highly AM-responsive, both at the phenotypic and transcriptomic level, suggesting that AM has great potential to promote argan growth under drought stress.

## Introduction

Drought is among the central threats associated with global warming and in many agricultural areas, it limits crop production. Prolonged drought imposes heavy metabolic costs on plants since it forces stomatal closure to reduce water loss, thereby limiting CO_2_ assimilation and photosynthetic efficiency (Díaz-Barradas et al. 2010). Furthermore, acute water stress triggers the production of reactive oxygen species (ROS) (Cruz de Carvalho 2008; Chakhchar et al. 2022) that interfere with many cellular processes. Plants mitigate the deleterious effects of drought stress through two distinct adaptive strategies that involve either stress avoidance or stress tolerance (Bandurska 2022; Levitt 1972). According to these strategies, plants can be categorized into two groups: the highly drought-adapted xerophytes, and the less specialized mesophytes, that are drought tolerant to various degrees, depending on their preformed stress avoidance and inducible stress tolerance mechanisms (Chen et al. 2024).

Extreme xerophytes in deserts can avoid drought stress during extended periods without water supply by extreme adaptations such as succulence (e.g. Cactuses), geophytism (Howard et al. 2019), or the ability to remain alive in a completely dried state (resurrection plants) (Marks et al. 2021). Less extreme xerophytes and drought-tolerant mesophytes exhibit constitutive adaptations such as a thick cuticle, high trichome density, or the separation of photosynthesis (during the day) from CO_2_ acquisition (at night) in the crassulacean acid metablism (CAM) (Heyduk 2022). We refer to these adaptations here as stress avoidance mechanisms, because they reduce the risk for plants to experience water stress.

In contrast to drought avoidance strategies, drought tolerance refers to inducible mechanisms that are triggered by drought stress to reduce the deleterious consequences of stress, once it has been perceived (Bandurska 2022). Drought tolerance involves physiological adaptations such as the accumulation of compatible solutes (sugars, amino acids, etc.) that replace water, and the induction of antioxidant pathways that mitigate oxidative damage caused by ROS (Bandurska 2022). Together, the drought-tolerance mechanisms protect cellular constituents and/or repair drought-related damage. Drought-coping mechanisms have been studied mainly in model species and crops that are amenable to physiological and genetic analysis. Perennial and woody species have received less attention, although they face unique challenges under extreme drought: Their extended vascular system is vulnerable to hydraulic failure (Li et al. 2022; Nour et al. 2024), and they cannot retract underground as geophytes, or survive as highly drought-resistant seeds as in annual plants (Liu et al. 2022). The argan tree (*Sideroxylon spinosum* syn. *Argania spinosa* L. Skeels) is considered a drought-adapted xerophyte (Terral et al. 2025), however, the underlying drought-resistance mechanisms are only poorly understood.

Many plant species profit from increased drought resistance conferred by beneficial associations with arbuscular mycorrhizal (AM) fungi (Formenti et al. 2026), and trees are no exception to that rule (Brunner et al. 2015; Liao et al. 2025). In particular their role in water balance and their beneficial effects on stomatal conductance and photosynthesis allow trees to better cope with drought stress (Augé et al. 2015; Smith and Read 2008). AMF also "prime" the host transcriptome, establishing a pre-alerted state that allows for a faster and stronger induction of ROS-scavenging mechanisms upon drought perception (Evelin et al. 2019). Argan is highly AM-responsive in terms of growth promotion (Nouaim and Chaussod 1994), and mitigation of water stress (Ganoudi et al. 2025). The recent availability of high-quality genomic resources in argan (Mateus et al. 2025; El Idrissi et al. 2026) provides a critical framework for studying the environmental adaptations of argan. Combined with emerging transcriptomic datasets (Rupp et al. 2024), these tools allow for a deeper understanding of the complex molecular cross-talk between abiotic stress and biotic interactions in argan.

Here, we use argan as an example of a xerophytic tree to study drought-coping strategies. Argan is endemic to the calcareous semi-arid and arid zones around the Moroccan cities of Essaouira, Agadir, Taroudant, and Tiznit. It constitutes the backbone of the ‘Arganeraie’, a unique ecosystem recognized as a UNESCO Biosphere Reserve in 1998 due to its critical biological, cultural, and socioeconomic values. Ecologically, *S. spinosum* acts as the final vegetative barrier against the northward encroachment of the Sahara desert (Msanda et al. 2005). Its deep and powerful root system plays an important role in stabilizing soils, maintaining hydraulic lift (Chakhchar et al. 2022; Tariq et al. 2026), and preventing erosion (Kirchhoff et al. 2019). Besides its ecosystem services, argan has a great socioeconomic impact with its high-value seed oil for cosmetic and culinary applications.

However, argan is under pressure by overuse as feed for goats and wood for heating and construction (Lybbert et al. 2011). In addition, the ‘Arganeraie’ faces an existential threat driven by the rapidly shifting climate of the Mediterranean and North African regions. A drastic reduction in precipitation, rising mean temperatures, and an increase in the frequency and severity of extreme drought episodes are predicted for the mediterranean basin (Lionello and Scarascia 2018). These intensifying arid conditions are threatening natural *S. spinosum* populations, by causing habitat fragmentation and range regression (Moukrim et al. 2019).

Apart from physiological responses to drought, we put the emphasis on the growth-promoting and protective effects of AM symbiosis for argan. While the general benefits of mycorrhizal symbiosis are well-established, they are highly context-dependent (Smith and Read 2008). The vast majority of experimental studies deal with the model symbiont *Rhizophagus irregularis* due to its global distribution (Benhiba et al. 2015; Essahibi et al. 2018; Savary et al. 2018) and the unique depth of genetic and genomic knowledge on this AMF (Tisserant et al. 2013; Manley et al. 2023). The generally low host specificity of AMF suggests that any fungal inoculum could potentially promote any AM-competent host plant, and this assumption is at the basis of many commercial applications (Chen et al. 2018a). However, the range of mycorrhizal growth effects varies greatly with the combination of AMF and host, resulting in some cases in pronounced growth depression (Klironomos 2003). These findings raise the question whether native fungal consortia, that have co-evolved with local hosts and with the local climatic and edaphic conditions, could confer superior mycorrhizal benefits than generic commercial inocula (Berruti et al. 2016; Essahibi et al. 2019).

Here we characterize the physiological response of argan to drought and to AM symbiosis, including ten native Moroccan inocula collected from the Arganeraie. Our results document the potential of argan to adapt to drought by a combination of preformed and inducible adaptive traits that promote stress avoidance and tolerance. In addition, we document that argan is highly AM-responsive, indicating the AM inocula may have great potential on restoration programs.

## Results

### Effects of drought and mycorrhizal status on argan growth

In order to identify AMF inocula with potential for growth promotion of argan and for protection against drought stress, we collected native inocula from 10 Moroccan sites in the Arganeraie region around Essaouira, Agadir, and Taroudant (**Fig. S1**). These were tested along with the drought-adapted AMF strain *Diversispora omaniana* isolated from a desert-like arid region of Oman (Symanczik et al. 2014), and with the established AM fungal model species *Rhizophagus irregularis* DAOM197198 (Tisserant et al. 2013). Plants were grown under well-watered conditions (80% field capacity (FC)), as well as under drought conditions (20% FC) for five months.

All inoculated plants exhibited AM colonization in a range of 35-95% colonization frequency) (**Fig. S2, Table S1**). In general, AM colonization was slightly lower in plants grown under drought, with one exception (inoculum S6) (**Fig. S2**). Global analysis of four major growth parameters (shoot height, shoot dry weight, root dry weight, and leaf area) showed two general trends over all treatments: drought severely reduced plant growth, and all inocula significantly increased it (**Fig. S3**). The strongest promotion of plant growth was achieved with inoculum S8 (**Fig. S3b,d**).

Prolonged serial measurements over 18 weeks confirmed that argan is highly AM-responsive (**Fig. S4**). Mycorrhizal plants grew at almost double the rate compared to non-mycorrhizal plants under well-watered conditions (**Fig. S4a**). Strikingly, non-mycorrhizal plants essentially ceased to grow under drought, while mycorrhizal plants continued to grow under drought (**Fig. S4b**), indicating that they have access to water and nutrients through the mycorrhizal network. These results show that AM symbiosis significantly stimulates the growth of argan independently of the water conditions, and that the negative consequences of drought can be compensated by AMF.

Total water uptake of plants was very low in all drought treatments irrespective of the mycorrhizal status (**Fig. S5a**). However, leaf relative water content (RWC) was maintained close to levels of well-watered plants (**Fig. S5b**). Consistent with an improved water status in mycorrhizal plants, stomatal conductance was significantly increased in mycorrhizal plants relative to non-mycorrhizal controls, both, in well-watered and drought-exposed plants (**Fig. S5c**). Inoculum S8 had accumulated the highest amounts of water under both irrigation regimes (**Fig. S5a**), consistent with its strong promotion of shoot growth (**Fig. S3b**).

### Global analysis of various physiological traits in response to drought and AMF

A multivariate analysis on all measured growth and physiological traits in plants inoculated with *R. irregularis* showed that the different samples formed four distinct clusters within the multidimensional space, indicating that both, water status and mycorrhizal colonization, highly influenced phenotypic variance (**Fig. S6**). We chose *R. irregularis* and three native inocula (S3, S8, S10) for global analysis of the influence of drought and mycorrhizal status on argan growth and physiology. Apart from the growth parameters discussed above (**Figs. S3, S5**), a series of traits related to drought response were determined: the number of shoot branches (SN), the levels of total soluble sugars and proline as compatible solutes (also known as osmolytes), four photosynthesis-related traits, namely chlorophyll content, non-photochemical quenching (PhiNPQ) as a measure of dissipated light energy, quantum yield of photosystem II (Phi2) as a measure of photosynthetic efficiency, and stomatal conductance, and, finally, four stress-related parameters, namely the fatty acid degradation marker malondialdehyde (MDA), the ROS hydrogen peroxide (H_2_O_2_), and the enzymes guaiacol peroxidase (POD) and catalase (CAT). Water deficit imposed a severe physiological cost on argan plants regardless of symbiotic status (with *R. irregularis*) (**Fig. 1**). On non-mycorrizal plants, drought significantly decreased growth, water uptake, and photosynthetic activity (lower Phi2; higher PhiNPQ), while at the same time elevating markers of oxidative stress (H_2_O_2_ and POD) (**Fig. 1**). When testing for the drought effect on mycorrhizal plants, we still observed a decrease in growth and water uptake, along with modification in photosynthetic activity and activation of oxydative stress and its mitigation (H_2_O_2_ and POD) (**Fig. 1**).

**Figure 1.**
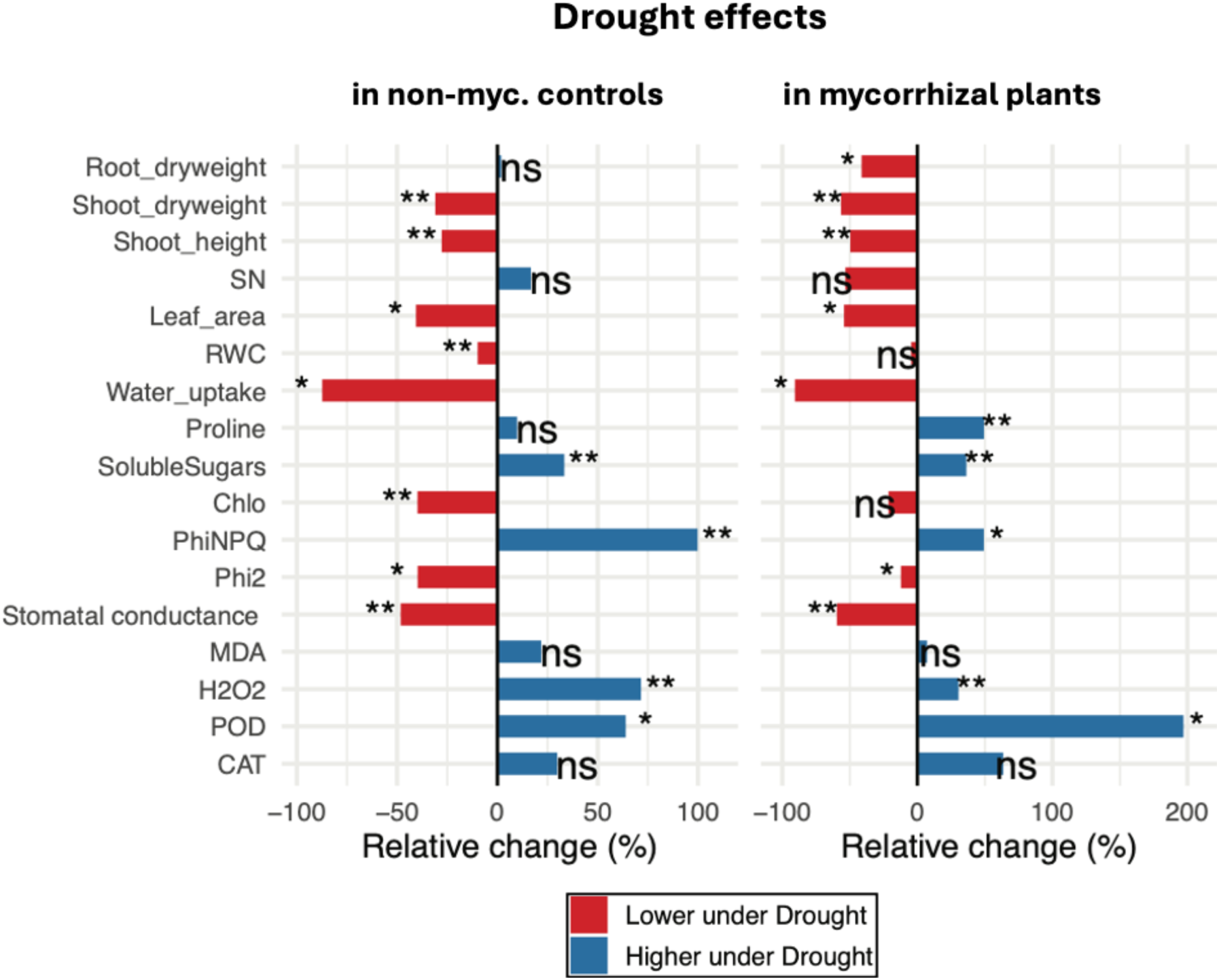
Effect of drought on various parameters related to plant growth and fitness. Relative difference (%) in 17 growth-related parameters determined in drought-treated plants vs. well-watered plants under control conditions, or in mycorrhizal plants (*R. irregularis* DAOM197198). The following parameters were determined: root dry weight, shoot dry weight, shoot height, number of lateral branches (SN), leaf area, relative water content of the leaves (RWC), accumulated water uptake during the experiment, the content of proline, soluble sugars, and chlorophyll (Chlo) in the leaves, non-photochemical quenching (PhiNPQ), quantum yield of photosystem II (Phi2), the levels of malondialdehyde (MDA) and hydrogen peroxide (H2O2, as well as the activities of (POD), and catalase (CAT). Bars represent mean values (n=5); significance levels (one-way ANOVA and Tukey’s test) are indicated as follows * *P* < 0.05; ** *P* < 0.01.

Mycorrhizal association with *R. irregularis* produced contrasting effects on argan. Under well-watered conditions, inoculated plants exhibited significantly higher biomass, water uptake and higher stomatal conductance compared to non-mycorrhizal controls (**Fig. 2**). Under drought stress, these symbiotic advantages on argan were maintained, although with an attenuated magnitude compared to well-watered conditions. Notably, mycorrhizal plants under drought displayed enhanced osmoregulation (soluble sugars, proline) and enhanced photosynthetic efficiency (lower PhiNPQ and higher Phi2) compared to non-mycorrhizal stressed plants (**Fig. 2**).

**Figure 2.**
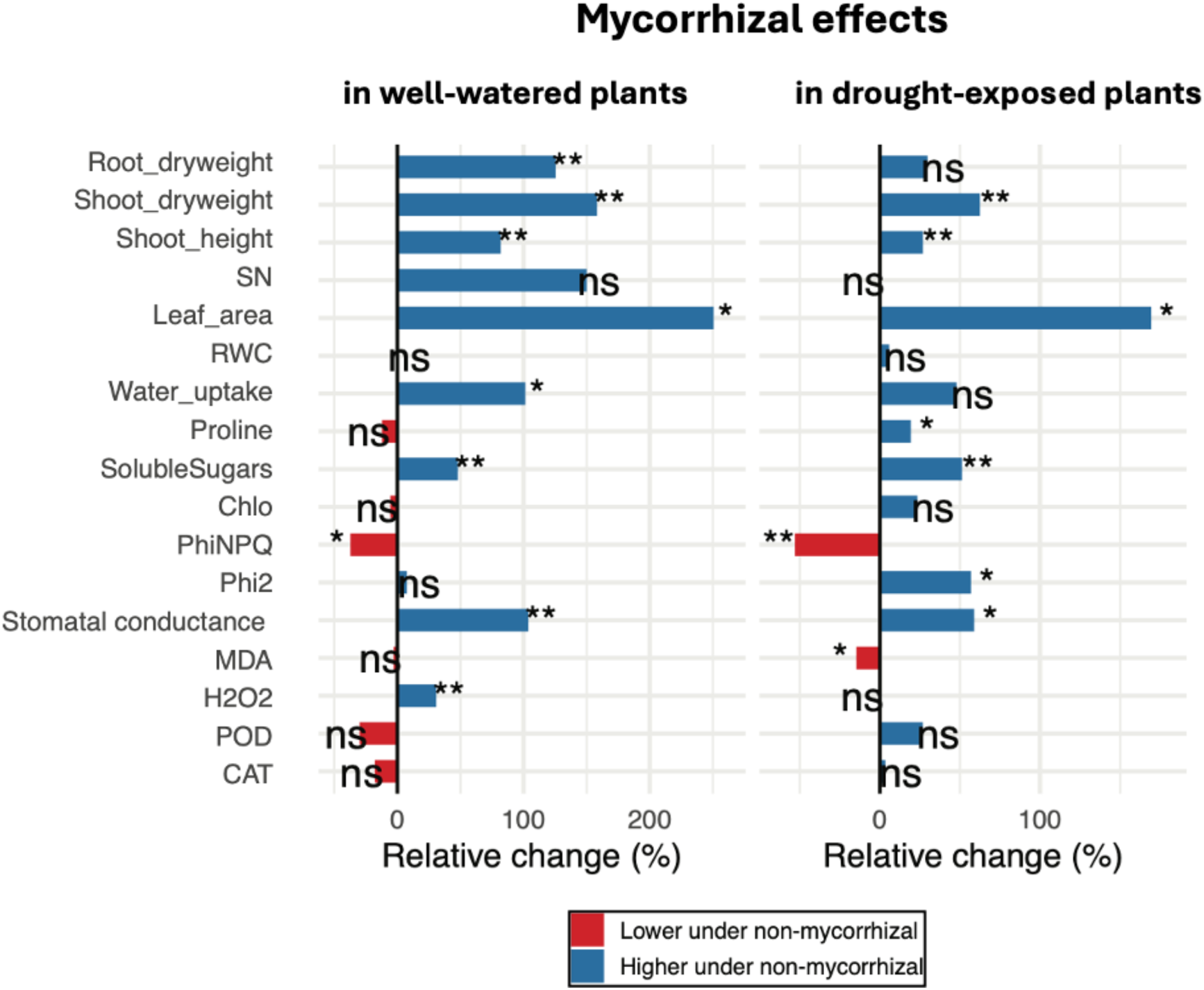
Mycorrhizal effects on various parameters related to plant growth and fitness. Relative difference (%) in 17 growth-related parameters determined in mycorrhizal plants vs. non-mycorrhizal controls under well-watered conditions or under drought. Eight weeks after inoculation with *R. irregularis* DAOM197198, the following parameters were determined: root dry weight, shoot dry weight, shoot height, number of lateral branches (SN), leaf area, relative water content of the leaves (RWC), accumulated water uptake during the experiment, the content of proline, soluble sugars, and chlorophyll (Chlo) in the leaves, non-photochemical quenching (PhiNPQ), quantum yield of photosystem II (Phi2), the levels of malondialdehyde (MDA) and hydrogen peroxide (H_2_O_2_), as well as the activities of (POD), and catalase (CAT). Bars represent mean values (n=5); significance levels (one-way ANOVA and Tukey’s test) are indicated as follows * *P* < 0.05; ** *P* < 0.01.

Next, we evaluated how the three native inocula (S3, S8, S10) affected global plant performance. In general, the native inocula had similar effects as *R. irregularis*, though to different degrees (**Fig. S7**). Under well-watered conditions, we observed a positive effect on plant growth, stomatal conductance, and several physiological parameters (**Fig. S7,** compare with **Fig. 2**). The beneficial effects of mycorrhiza on plant growth and photosynthetic parameters were also observed under drought (**Fig. S8,** compare with **Fig. 2**), although the mycorrhizal benefit on plant growth was weaker than under well-watered conditions (compare **Fig. S8** with **Fig. S7**). Importantly, AM under drought increased protection against oxidative damage, as exemplified by lower MDA levels and increased catalase activity (**Fig. S8**). In the comparison of the three native inocula, **S8** displayed the strongest beneficial effects (**Fig. S7, S8**), also relative to *R. irregularis* (compare with **Fig. 1**), particularly in mitigating drought-induced biomass loss, and in promoting shoot branch numbers (SN). Global analysis by clustering between well-watered and drought-stressed plants with the four inocula (*R. irregularis*, S3, S8, S10) confirmed the dominant effect of water regime, and the strong growth-promoting effect of S8 under both water regimes (**Fig. S9**).

While drought had strong effects on growth parameters (**Figs. 1,2; Figs S3, S4, S7, S8**), the ROS intermediate H_2_O_2_ showed only moderate drought-related induction of approximately 30% and 75% in mycorrhizal and non-mycorrhizal plants, respectively, and the membrane lipid oxidation marker MDA was not affected in either case (**Fig. 1**). To detect potential local accumulation of H_2_O_2_ in drought-stressed plants, we performed DAB staining in leaves as described (Daudi and O’Brien 2012). Surprisingly, no induction was observed in drought-stressed plants (**Fig. S10**), indicating that the levels of ROS in drought-stressed argan leaves are below the sensitivity threshold of this technique. As a control for the effectivity of the assay, we subjected maize plants to a similar treatment and assessed shoot growth and leaf H_2_O_2_ levels in the same way as in argan. This experiment revealed strong growth defects and H_2_O_2_ accumulation in stressed maize leaves (**Fig. S11**), validating the employed H_2_O_2_ assay.

### Transcriptomic response of argan to drought and mycorrhizal colonization

To explore the effect of drought and mycorrhizal colonization on gene expression patterns in argan, we performed an RNA sequencing (RNAseq) experiment. The most beneficial native inoculum (S8) was employed for this analysis. A factorial experimental setting with drought and inoculum S8 with the respective controls was performed with three independent replicates each, resulting in a total of 12 samples. We obtained on average 33 million reads per sample (**Supplementary Table S2**). While the drought-treated replicates (20% FC) generally clustered tightly in PCA analysis, the well-watered samples (80% FC) exhibited a wider distribution in particular across the PC1 axis, which explained 47 % of variance (**Fig. S12**).

As a first step, we used the sequence data to characterize the mycorrhizal species in the inoculum used for RNAseq. First, the RNAseq reads were pseudo-mapped in parallel to different AMF species representing several AMF families. The multiple mapping analysis allowed us to obtain a proxy for AMF composition in the mycorrhizal RNAseq samples. As expected, non-mycorrhizal samples did not contain any mycorrhizal reads, while the native inoculum (S8) contained a mixture of several members of different AMF genera including *Funneliformis*, *Rhizophagus*, *Glomus*, *Claroideoglomus* and *Paraglomus* (**Fig. S13**). While most of the samples exhibited a similar fungal composition, the sample 1A showed a distinct AMF pattern, indicating that the initial AMF consortium isolated from Oulad Teima (**Fig. S1**) had segregated atypically in this sample, potentially explaining the wider spreading of the well-watered mycorrhizal samples in PCA analysis (**Fig. S12**).

### Gene expression patterns in argan roots exposed to drought and AMF

Next, we evaluated the global transcriptional changes driven by water status (drought vs. well-watered) and symbiotic status (mycorrhizal vs. non-mycorrhizal) (**Supplementary Table S3**). First, the drought effect on the argan transcriptome was examined. In non-mycorrhizal plants, drought triggered a moderate transcriptional shift, resulting in 810 differentially expressed genes (DEGs) (**Fig. 3a**). Among the top 0.5% up-regulated genes, were two heat shock proteins (HSP70) (Usman et al. 2017), a stress-related lipid transfer protein (LTP2) (Salminen et al. 2016), and a marker of ABA-dependent responses (AWPM-19) (Yao et al. 2018) (**Fig. S14**).

**Figure 3.**
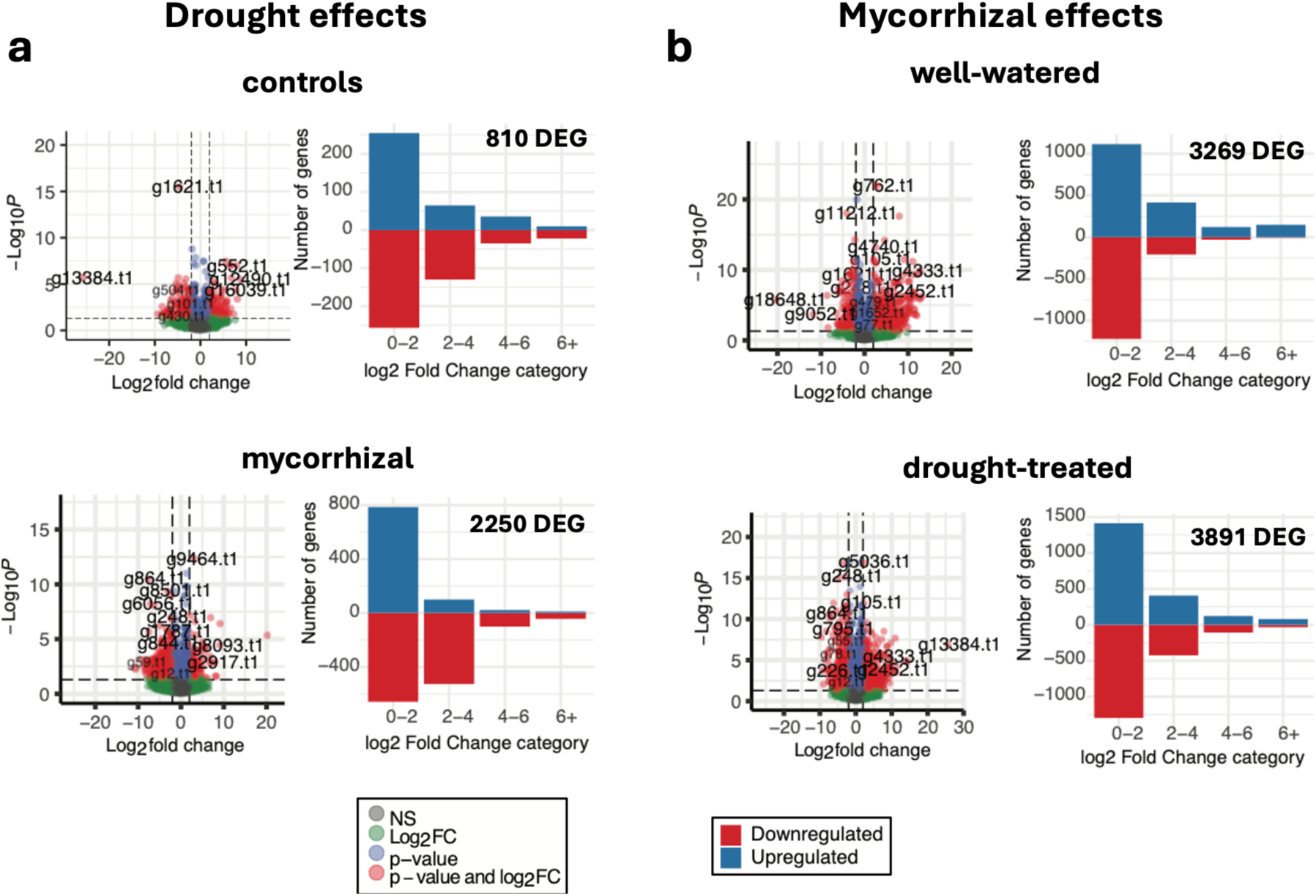
Effects of drought and mycorrhizal status on gene expression in argan. Differentially expressed genes are shown in volcano plots with indication of the fold change (x-axis; log2) and the significance level for regulation (y-axis; -log10). Histograms show the different log2Fold change categories of differentially expressed genes as indicated **a**) Drought effect in non-mycorrhizal plants (drought vs. well-watered in non-mycorrhizal controls) and drought effect in mycorrhizal plants (drought vs. well-watered in mycorrhizal plants). **b**) Mycorrhizal effect in well-watered plants (Mycorrhiza vs. non-mycorrhiza in well-watered plants) and mycorrhizal effect in drought-treated plants (Mycorrhiza vs. non-mycorrhiza in drought).

In mycorrhizal plants, the drought-induced transcriptomic response was substantially larger with 2250 DEG (**Fig. 3a**), of which 179 were shared with drought-responsive genes in control plants (**Fig. 4a**). Among the top drought-regulated genes in mycorrhizal roots, we observed the stress response protein thaumatin (Park and Kim 2021), metallothionein (Saeed Ur et al. 2020), and again LTP2 (**Fig. S15**).

**Figure 4.**
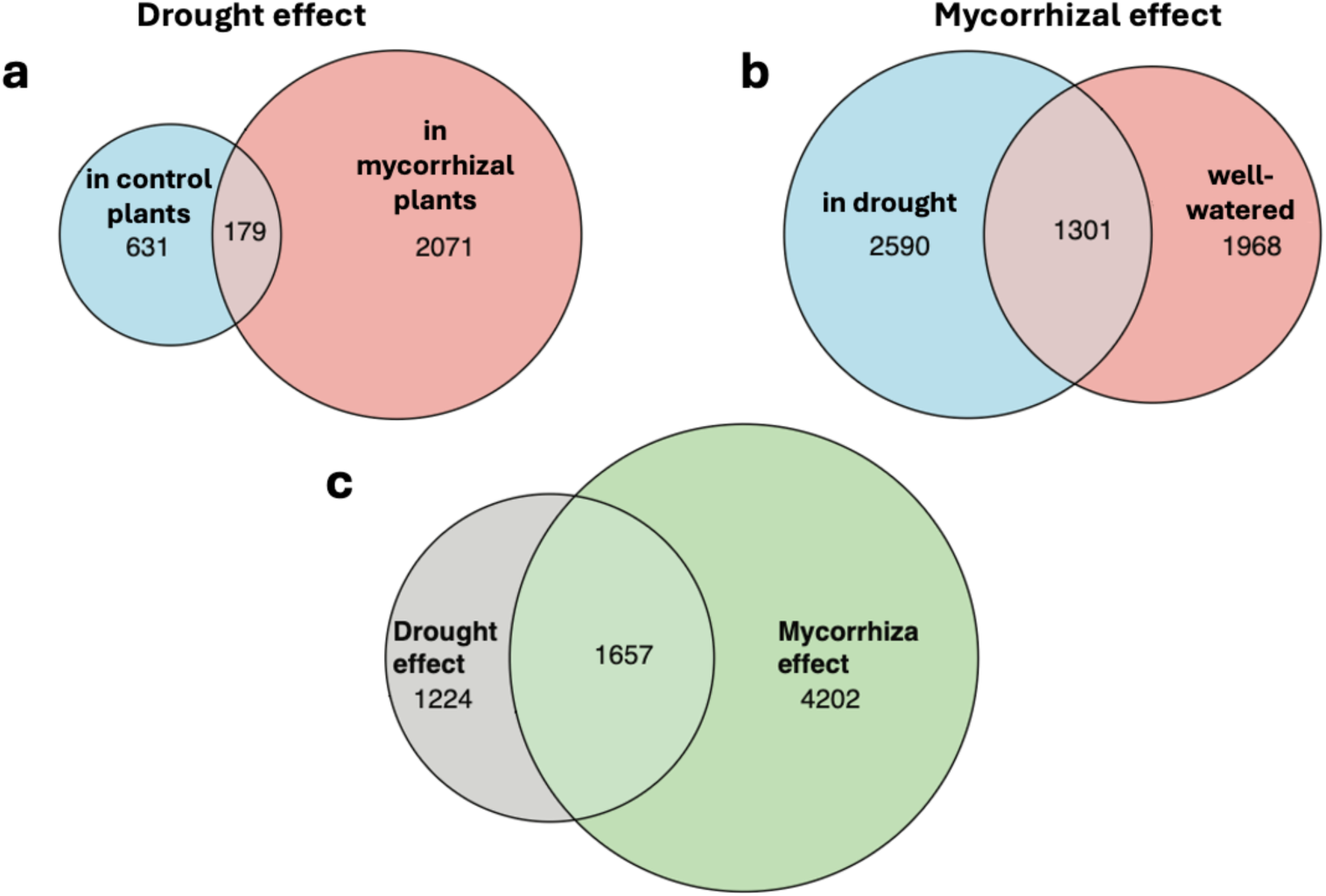
Global gene expression patterns related to drought and mycorrhizal status. Venn diagrams showing differentially expressed genes shared betweeen the different conditions. **a)** Drougth-responsive genes shared between mycorrhizal and non-mycorrhizal conditions. **b**) Mycorrhiza-responsive genes shared between plants under drought and under well-watered conditions. **c**) Summary analysis showing the number of genes overlapping between the drought-responsive and AM-responsive genes.

The mycorrhizal response was generally more pronounced than the drought response with 3269 genes differentially expressed in well-watered conditions and 3891 AM-responsive genes under drought (**Fig. 3b**), including 1301 shared genes (**Fig. 4b**). AM responsive genes under well-watered conditions included an induced chitin-binding protein that could promote symbiosis by sequestering immunogenic chitin oligomers (Bakhat et al. 2023) (**Fig. S16**). In addition, several known AM-inducible genes were observed, such as GST and peroxidase (Breuillin et al. 2010) (**Fig. S16, S17**).

In general, we observed that the mycorrhizal status regulated a significantly larger unique gene set (4202 unique DEGs) compared to the water status (1224 unique DEGs), with a core set of 1657 DEGs responding to both drivers (**Fig. 4c**).

To systematically address the biological processes affected by these transcriptional shifts, we performed a Gene Ontology (GO) enrichment analysis visualized according to neighbourhood in a semantic space (Reijnders and Waterhouse 2021) (**Fig. S18**). This approach revealed that the drought response was characterized by the enrichment of structural and signaling pathways, specifically “plant-type secondary cell wall biogenesis”, “auxin-activated signaling pathway” and “polysaccharide metabolic process”. In contrast, the main mycorrhizal effect displayed a distinct functional signature dominated by metabolic and defense-related processes. Key enriched terms included "hormone metabolic process", "terpenoid metabolic process" and "response to lipid".

To assess the transcriptional response of AM-related core genes, we first identified the argan orthologs of well-established AM-related marker genes (Bravo et al. 2016; Favre et al. 2014) (**Table S4**) and assessed their differential expression between inoculated and non-inoculated plants in well-watered and drought stress conditions. Genes related to the common symbiosis signaling pathway (CSSP), and to mycorrhizal infection showed weak responsiveness to mycorhizal status, with a tendency to downregulation under drought (**Fig. 5**). In contrast, homologs of genes expressed in cells with arbuscules and involved in symbiotic nutrient exchange (*RAM1*, *RAM2*, *STR*, *PT4*, and *AMT2*) were induced in mycorrhizal roots relative to non-mycorrhizal roots, with the exception of STR2 (**Fig. 5**). Interestingly, drought slightly attenuated mycorrhizal induction of arbuscule-related genes, in particular the induction of the transcriptional activator of many AM-related genes, *RAM1* (Gobbato et al. 2013; Pimprikar et al. 2016; Rich et al. 2017), was reduced by drought (**Fig. 5**).

**Figure 5.**
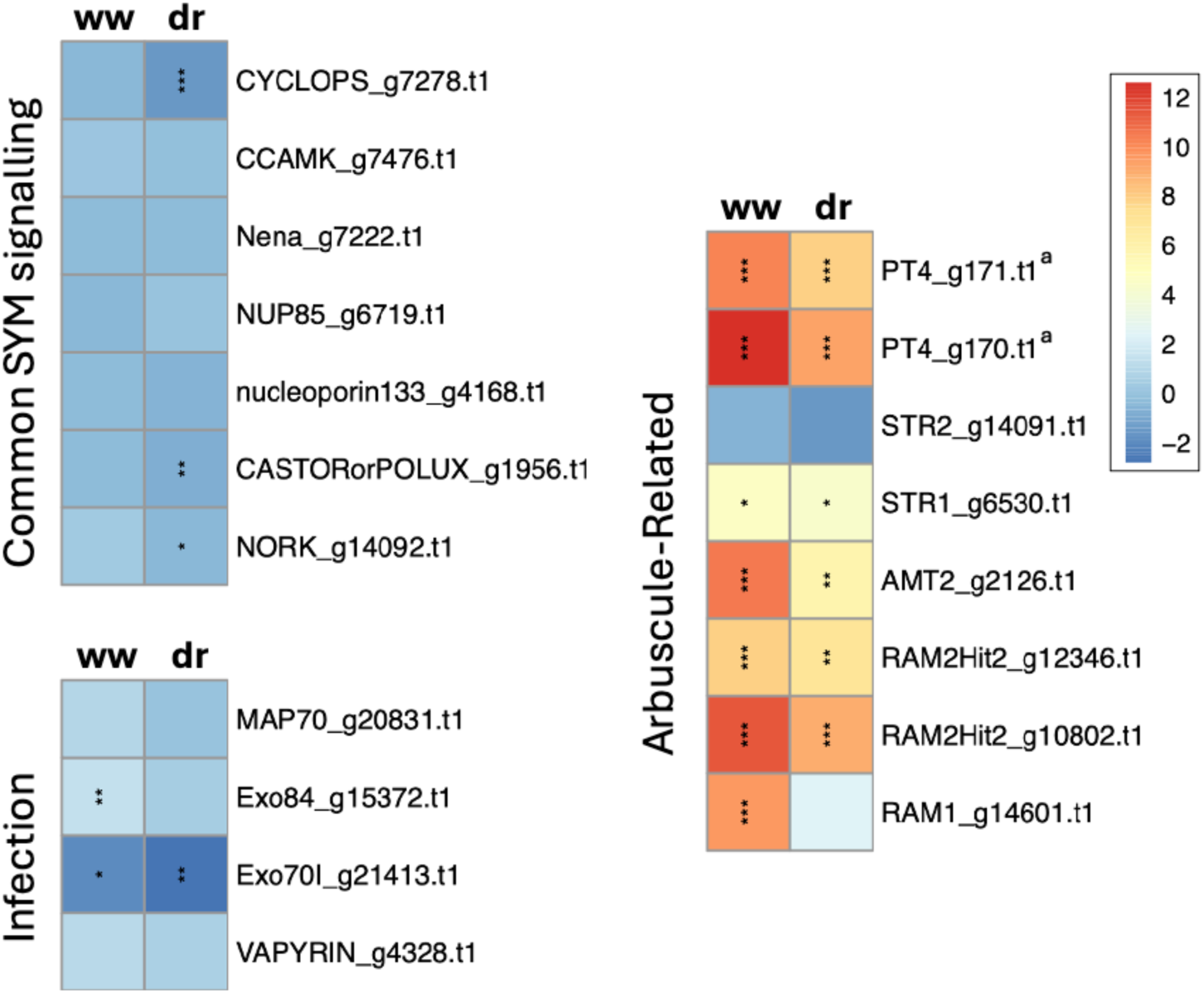
Expression pattern of argan orthologues of conserved mycorrhiza-related genes. First, the closest argan homologues of conserved mycorrhiza-related genes according to Bravo et al., and Favre et al., were identified. Their induction in mycorrhizal roots relative to non-mycorrhizal controls is shown for well-watered conditions (WW) and for drought-treated plants (DR). Genes were grouped according to their functional importance in real signaling (common SYM signaling), root infection (infection), or their importance for bidirectional nutrient exchange in host cells with arbuscules (arbuscule-related). The heatmaps show fold change (log2) in 3 independent samples per treatment. ^a^ The two PT4 orthologs were allocated in a tandem duplicated arrangement in haplotig 2 (Mateus et al., 2025). Significance levels show adjusted p-values between non-mycorrhizal and mycorrhizal plants from the DEseq2 anaylsis. Adjusted p-values are indicated as follows * *P* < 0.05; ** *P* < 0.01 and *** *P* < 0.001.

To specifically identify argan genes relevant for stress resistance, we compiled a curated list of genes assigned to the two major strategies, drought avoidance and drought tolerance, based on a comparative analysis of drought-related genes in *Arabidopsis thaliana* and the xerophyte *Reaumuria soongorica* (Shi et al. 2013). Next, we identified the argan orthologs of Arabidopsis genes related to drought avoidance and drought tolerance, respectively (**Supplementary Table S5**), and analyzed their expression patterns. Surprisingly, most of the genes did not show a significant response to drought, neither in mycorrhizal, nor in non-mycorrhizal roots (**Fig. 6**). Among a total of 88 genes assigned to drought response, only 7 and 11 genes (8% and 12.5%) were drought-responsive in non-mycorrhizal and mycorrhizal roots, respectively (**Fig. 6**).

**Figure 6.**
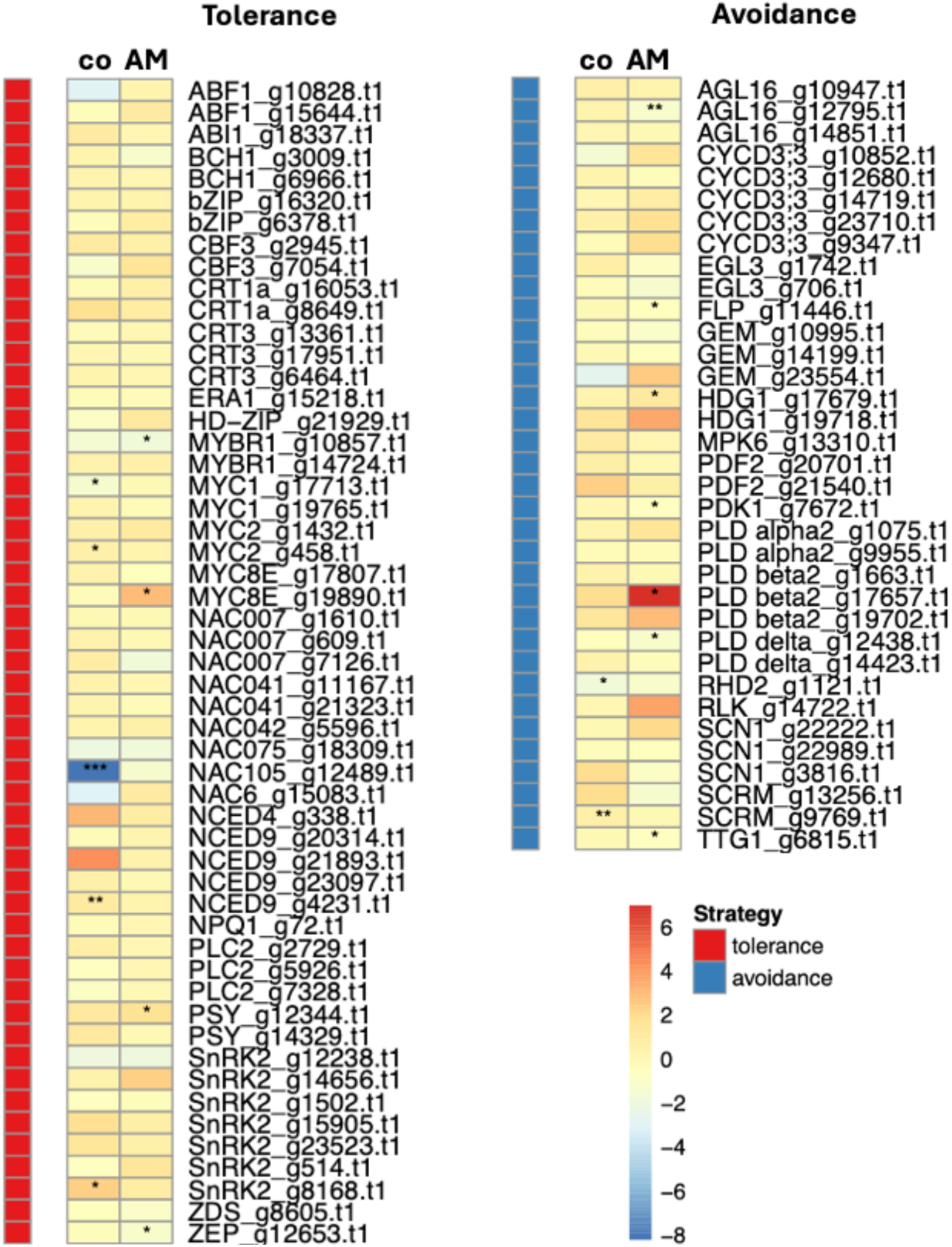
Expression pattern of drought-related genes. First, the closest argan homologues of drought-related genes were identified according to genes assigned to drought avoidance due to their role in preformed adaptations (cuticle, trichomes etc.), or drought tolerance based on their predicted role in inducible drought resistance (ABA-related genes). Drought-induced changes in gene expression of the closest argan homologues is shown in mycorrhizal plants (AM) and in non-mycorrhizal controls for the two categories drought stress tolerance (left) and drought stress avoidance (right). The heatmaps show fold change (log2) in 3 independent samples per treatment. Significance levels show adjusted p-values between well-watered and drought plants from the DEseq2 anaylsis. Adjusted p-values are indicated as follows * *P* < 0.05; ** *P* < 0.01 and *** *P* < 0.001.

## Discussion

Argan is a well-adapted xerophyte protected from drought by constitutive stress-avoidance traits and inducible stress-tolerance mechanisms (Barradas et al. 2013; Chakhchar et al. 2022; Chakhchar et al. 2017; Chakhchar et al. 2018; Chakhchar et al. 2024; Terral et al. 2025; Timzioura et al. 2025). Yet, extension of the drought period, and a generally warmer climate threaten the Arganeraie in Morocco. Here, we describe the drought response of argan at the physiological and transcriptomic level, and we explore the potential of arbuscular mycorrhiza (AM) to protect argan against drought. A better understanding of the molecular drought adaptation mechanisms in argan will help protect the Arganeraie against overuse and progressive decline (Lybbert et al. 2011), and it will enable further development of the unique species by marker-assisted breeding, based on the recent description of high-resolution chromosome-level genome sequences for argan (El Idrissi et al. 2026; Mateus et al. 2025).

Argan is highly AM-responsive (**Fig. S3**), in particular under well-watered conditions, i.e. it grows larger and produces more foliage. Mycorrhizal effects were also observed under drought, but the degree of growth promotion depended on the inoculum. The beneficial effects on stomatal conductance and relative water content suggests that AM allows plants to acquire more CO_2_ while limiting water loss through stomates. This may contribute to overall reduced stress levels, while growth can be sustained longer than in non-mycorrhizal plants.

Notably, H_2_O_2_ levels were only moderately increased in argan leaves under drought (31% induction in AM; 72% in controls), compared to increases of, for example, 125-140% in the more drought-sensitive wheat (Ostrowska et al. 2024). In general, H_2_O_2_ levels were relatively low with a range between 0.11 and 0.38 µM, compared to steady-state concentrations between 0.67 and 3.63 µM in the leaves of seven angiosperms with different life styles (Cheeseman 2006). Consistent with low levels of oxidative stress, the membrane oxidation marker MDA was not affected in drought-treated argan leaves, suggesting that argan leaves under drought are protected from oxidative damage by stress avoidance and stress tolerance mechanisms.

In line with this notion, the argan transcriptome showed a weaker response to drought than to AM, and a selection of drought-related genes, identified by their role in stress avoidance and stress tolerance under conditions of drought in Arabidopsis and the desert plant *R. soongorica* (Shi et al. 2013), was only moderately affected by drought (**Fig. 6**). The weak (non-significant) induction of the H_2_O_2_-converting enzyme catalase is in line with low oxidative stress levels.

Taken together, these results suggest that argan has stress-avoidance mechanisms that prevent a level of stress that would be associated with increased levels of ROS markers such as H_2_O_2_ and MDA. Nevertheless, argan has undoubtably also a wide range of inducible drought tolerance mechanisms, perhaps the most extreme reaction being the shedding of all its leaves under extreme drought to minimize water loss (Terral et al. 2025), a strategy that allows prolonged survival at a limited metabolic cost.

Future work will systematically address the identity of native beneficial inocula to tailor optimal microbial communities to protect argan from environmental stress. All our inocula contained AM fungi, and the overall similarity of the effects of native Moroccan inocula with the effects of *R. irregularis* and *D. omaniana* testify for the potential of AM fungi to protect argan. Nevertheless, a contribution of other members of the argan microbiome, including plant-growth-promoting rhizobacteria (PGPR), cannot be excluded. PGPRs can promote plant growth, improve mineral nutrition, and increase resistance against pathogens (Lugtenberg and Kamilova 2009). The mechanisms by which PGPR improve plant fitness is a combination of solubilization of soil nutrients, modulation of hormonal homeostasis, and induction of induced systemic resistance (ISR) (Lugtenberg and Kamilova 2009; Pieterse 2025). PGPR are ubiquitous and can protect crops against drought (Chieb and Gachomo 2023), however, their contribution to plant growth promotion is likely to be limited in highly arid conditions such as the Arganeraie during the summer season. AM fungal mycelia can acquire residual water from microscopic cracks in soil particles and from deep soil, thereby promoting plant fitness even under extreme desert conditions (Etesami et al. 2026; Madouh et al. 2025; Vasar et al. 2021). Bacteria, on the other hand, are more sensitive to drought than fungi and often enter a resting state under extreme drought (Metze et al. 2023; Schimel 2018). Also, plants secrete lower amounts of exudates to rhizobacterial communities (Fuchslueger et al. 2014), hence PGPRs are less likely to confer beneficial effects to argan than AM fungi during the dry season. Nevertheless, complex synthetic communities encompassing fungal and prokaryotic microbes should in the future be evaluated for their protective effects on argan.

Based on our results and previously published work, we conclude that argan is well-adapted to survival under prolonged drought, using a combination of stress-avoidance and stress-tolerance strategies. In addition to these endogenous adaptive traits, AM have great potential to protect argan from severe drought. These features will be essential to cope with increasing temperatures and extended drought periods associated with global warming.

## Methods

### Physiological experiments

#### Biological material and experimental design

Mature seeds of argan were disinfected with 5 % bleach solution, rinsed and submerged in sterile distilled water for 10 days. Treated seeds were then sown in sterile mixture of sand and soil (3:1) and kept for two months in a growth chamber under 25/22 °C Day/night temperature, 60% relative humidity, and a photoperiod of 16 h. The most similar seedlings were selected and transferred into plastic pots containing 1.2 kg of the same sterile substrate used for germination. At transplanting, plants were divided into non-inoculated plants (NM), plants inoculated with *Rhizophagus irregularis* (DAOM197198) and the drought-adapted AM fungus *Diversispora omaniana* (Symanczik et al. 2014) and plants inoculated with field inocula collected from the rhizosphere of argan trees in natural populations at Morocco around Essaouira, Agadir, and Taroudant. The inoculum used in these experiments consisted of 10 g of rhizospheric soil containing spores, extra-radical mycelium, and infected roots (chive for RI and argan for native inoculi). The control plants (NM) received an equal quantity of autoclaved inoculum. After transplantation, plants were well watered to 80% Field Capacity (FC: the maximal water retention capacity) for 3 months, and then each group was divided into two categories. The first category was kept well-watered to 80% FC (Well-watered) and the other category was exposed to severe water stress by reducing the water regime to 20% FC (Drought). To keep each water regime, the pots were examined by weighing and the amount of water lost was replaced. The physiological experiments were carried out in the same growth chamber with the same conditions as described above

### Mycorrhizal colonization and plant growth parameters

After 2 months of water deficit application, plants height and shoots number were measured. The plants were then harvested and the shoots and roots were separated and dried at 70 °C for 48 h to measure shoot and root dry weights. Mycorrhizal status of the plants was assessed as described (Phillips and Hayman 1970) with some modifications. At harvest, fresh roots were collected, thoroughly washed with distilled water, and cleared with 10% KOH at 90 °C for 2 h. The roots simples were then washed, treated with 7.5% H2O2 for 10 min, and then acidified in 1% HCl for 10 min. The prepared roots were stained with 0.05% (w/v) trypan blue in lactoglycerol at 90 °C for 20 min. The stained roots were then cut into 1 cm pieces and observed under an optical microscope. The frequency and intensity of roots colonization were calculated as described (Trouvelot et al. 1986). For determination of leaf area, digital photographs of leaves were analyzed using ImageJ software (https://imagej.net/ij/download.html).

### Relative water content (RWC), and water uptake (WU)

Fresh leaves were harvested, weighted and then submerged in water for 24 h at 4 °C in darkness to determine turgid weight after fully hydrating. The leaves were then dried at 80 °C for 48 h to measure dry weights. RWC was determine using the following formula (1) (Talaat and Shawky 2014):

RWC = ((FW − DW)/ (TW − DW)) x 100, where FW, DW and TW are fresh, dry and turgid weigh, respectively.

Water uptake was determined by calculating the total volume of water added during the period of the experiment minus the volume of water evaporated. During the experimentation period, pots were weighed every two days and adjusted to a predefined weight (weight of the pot containing substrate, plant, and the amount of water corresponding to each water regime) by adding water. The volume of water added to maintain this predefined weight subtracted from the volume of water evaporated gives the amount of water absorbed by the plant during each weighing (Meddich et al. 2000). The amount of water evaporated was measured using the same method with pots containing the same substrate and conditions but without the plant.

### Stomatal conductance and chlorophyll fluorescence

Stomatal conductance was measured using a photosynthesis system (LI-6400/XT, Version 6). The chlorophyll fluorescence was measured by using a portable

fluorometer (MultispeQ, V 2.0).

### Determination of total chlorophyll content

Total chlorophyll content was determined as described (Arnon 1949). Fresh leaves (100 mg) were ground in a cold mortar with 2 mL of acetone (80%, v/v) and centrifuged at 5,000 *g* for 10 min. Chlorophyll content was determined by measuring the absorbance of the supernatant at 663 and 645 nm and calculated using the following formula:

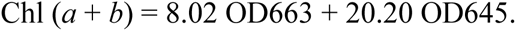

### Determination of total soluble sugars (TSS) and proline content

TSS were extracted from fresh leaf tissues (100 mg) in potassium phosphate buffer (50 mM, pH 7.5). TSS contents were measured according to (Irigoyen et al. 1992) by mixing 0.1 mL of the extract with 3 mL of freshly prepared anthrone (200 mg anthrone, 100 mL 72% H2SO4). The reaction mixtures were boiled for 10 min, cooled, and then the absorbance was measured at 620 nm. The concentrations of TSS were calculated using a glucose standard curve.

Proline extraction and determination was also performed in extracts of fresh tissue (100 mg) using the ninhydrin method according to (Bates et al. 1973). Proline content was measured by spectrophotometric analysis at 515 nm and calculated using a proline standard curve.

### Determination of malonyldialdehyde (MDA) and hydrogen peroxide (H_2_O_2_) content

Lipid peroxidation was assessed by determination of MDA content as described (Hernández and Almansa 2002). Fresh leaves (100 mg) were ground in a cold mortar in 2 mL of 0.1% (w/v) trichloroacetic acid (TCA). After centrifugation of the homogenate at 14,000 g for 10 min, the supernatant (0.5 mL) was mixed with 1.5 mL of TCA at 20% containing 0.5% of thiobarbituric acid (TBA). The mixture was heated at 90°C for 20 min, cooled quickly on ice, and centrifuged at 10,000 g for 5 min. The absorbance of the supernatant was measured at 532 nm and at 600 nm for the correction of nonspecific turbidity corrected by (A600) subtracting from (A532). MDA concentrations were calculated using the molar extinction coefficient (155 mM−1 cm−1).

Hydrogen peroxide concentrations were determined according to (Velikova et al. 2000), with some modifications. Fresh leaves (100 mg) were homogenated in a cold mortar with 2 mL of 20 % (w/v) trichloroacetic acid (TCA), centrifuged for 15 min at 12,000*g* in 4 °C. The supernatant (0.5 mL) was mixed with 0.5 mL potassium phosphate buffer (10 mM, pH 7.0) and 1 mL of 1 M potassium iodine (KI). The reaction mixture was incubated for 1 h in the dark, and then the absorbance was determined at 390 nm. H_2_O_2_ concentrations were determined in reference to a standard curve established with known concentrations of H_2_O_2_.

### Histochemical hydrogen peroxide (H_2_O_2_) detection

H_2_O_2_ was detected using DAB (Diaminobenzidine) according to the method described by Vanacker et al (2000). Fresh leaves were collected and immersed in DAB buffer, and then incubated overnight in the dark to allow DAB uptake and reaction with H_2_O_2_ and peroxidase. DAB polymerizes locally as soon as it comes into contact with H_2_O_2_ in the presence of peroxidase, giving a reddish-brown polymer. The prepared leaves were decolorized in ethanol at 90 °C for 30 min and then observed under a microscope.

### Antioxidant enzyme essays

Fresh leaves (200 mg) were grounded into a fine powder with liquid nitrogen and homogenized in 2 mL of potassium phosphate buffer (0.1 M, pH 7), 1 mM EDTA, and 1% PVPP (polyvinylpolypyrrolidone). The homogenate was centrifuged at 15,000 *g* at 4 °C for 15 min and the supernatant is used for subsequent enzymatic assays.

Catalase (CAT) activity was evaluated following the method of (Aebi 1984) using H_2_O_2_ as substrate. The assay mixture consisted of 0.1 M phosphate buffer (pH 7.0) contains 10 mM H2O2 and 100 μL enzyme extract. The absorbance was recorded at 240 nm for 3 min and an extinction coefficient (E) = 39.4 mM−1 cm−1 was used to calculate CAT activity. Guaiacol peroxidase (POD) activity was assayed as described (Hori et al. 1997). The reaction mixture consisted of enzyme extract (0.1 mL), 50 mM phosphate buffer (pH 7.0), 40 mM guaiacol, and 10 mM H_2_O_2_. The absorbance was recorded at 470 nm for 3 min and the activity was determined using an extinction coefficient of 25.5 mM−1 cm−1.

### Statistical Analysis

All statistical analysis were performed using R software (Team). The effects of mycorrhizal inoculation and drought stress on argan physiology were statistically analyzed with ANOVA, where mycorrhizal status and water-status were used as first and second factor.

### RNA Sequencing and Bioinformatics Analysis

The effects of drought and mycorrhizal colonization were assessed by RNA-sequencing. The experimental design consisted of four different treatments: well-watered, drought, mycorrhizal (inoculum S8), and non-mycorrhizal (inoculated with autoclaved inoculum) with three replicates each. Plants were grown for 5 months in a growth chamber with their respective treatments. Fine roots were sampled, selected for extraradical fungal colonization, and snap frozen. RNA extraction was performed by BGI Tech Solutions as follows: The appropriate amount of tissue was ground into powder under liquid nitrogen and transferred to 1.5ml of preheated CTAB lysis reagent (add 2% β-mercaptoethanol) at 65°C, and incubated at 65°C for 15 minutes. After incubation, the tube was cooled to room temperature, then centrifuged at 4°C with 12,000xg for 5 minutes. The supernatant was transferred to 2.0ml EP tubes. 200μL chloroform/isoamyl alcohol (24:1) per ml of CTAB lysis buffer was added. The mixture was vortexed to mix and centrifuged at 4°C with 12,000xg for 10 minutes.

Next, the clear upper aqueous layer was transferred to a 2.0ml EP tube. An Equal volume of phenol/chloroform/isoamyl alcohol (25:24:1) was added. The mixture was vortexed to mix and centrifuged at 4°C with 12,000xg for 10 minutes. Then the clear upper aqueous layer was transferred to a 2.0ml EP tube. An equal volume of chloroform/isoamyl alcohol (24:1) was added. The mixture was vortexed to mix and centrifuged at 4°C with 12,000xg for 10 minutes. The final clear upper aqueous layer was transferred to a new 1.5mL EP tube and 2/3 the volume of the aqueous phase of isopropanol was added. The mixture was gently inverted to mix well and placed at -20°C for 2 hours. Precipitated RNA was collected through centrifugation at 4°C with 12,000×g for 20 minutes. The supernatant was removed, and the pellet was washed with 1ml of 75% ethanol and dried in the biosafety cabinet for 3-5 minutes. Finally, 20µL∼200µL of DEPC-treated RNase-free water was added to dissolve the RNA. Total RNA was qualified and quantified using a Agilent 2100 Bioanalyzer (Agilent, CA, USA) before library preparation with the DNBSEQ stranded mRNA library protocol. Sequencing libraries including poly-A selection, were constructed using the MGIEasy RNA Library Prep kit according to the manufacturer’s protocol. The resulting libraries were sequenced on an DNBseq platform to generate 100 bp paired-end reads. All raw sequence data were submitted to the EBI/ENA repository (accession Project No.: PRJEB60382). Raw sequencing data underwent quality control and preprocessing using fastp (Chen et al. 2018b) to remove adapter sequences and trim low-quality bases. High-quality reads were then aligned to the *Sideroxylon spinosum* haplotype 1 genome assembly (Mateus *et al.,* 2025) with a two-pass mapping strategy using STAR v2.7.10 (Dobin et al. 2013). Post-alignment, gene-level abundance was quantified by counting read fragments assigned to genomic features using featureCounts (Liao et al. 2014).

Differential expression analysis was performed in the R statistical environment using the DESeq2 Bioconductor package (Love et al. 2014). The raw count matrix was imported and converted into a DESeqDataSet object using the DESeqDataSetFromMatrix function. To reduce noise and improve statistical power, the dataset was pre-filtered to remove low-abundance transcripts, retaining only features with a total sum of at least 10 counts across all samples. Differential expression was tested using a negative binomial generalized linear model (GLM). Specific contrasts were applied to evaluate the effects of drought (Drought vs. well-watered) and mycorrhizal colonization (AM vs. NM), as well as their interaction. P-values were adjusted for multiple testing using the Benjamini-Hochberg procedure. Enriched GO terms were identified with the R package topGO (Alexa et al. 2006). The different gene features were functionally annotated with eggNOG (Cantalapiedra et al. 2021). The gene universe is defined as all annotated argan genes and the genes of interest are defined as the differentially expressed genes in each comparison. We visualized the different enrichment results on a semantic space with the Go-figure tool (Reijnders and Waterhouse 2021) with default commands. All scripts and code used for this analysis are available in the GitHub repository: https://github.com/ivandamg/Argan_RNAseq.

### Assessment of the mycorrhizal composition of inoculum S8

The composition of the AM fungal community in inoculum S8 was characterized directly in colonized argan fine roots sampled for RNA-Seq analysis. We developed an *in silico* taxonomic profiling pipeline. Following quality control and adapter trimming with fastp (Chen et al. 2018b), high-quality reads were aligned to a custom reference database comprising representative genomes from the major classes of the Glomeromycotina phylum (Montoliu-Nerin et al. 2021). Transcript abundance for each fungal species was quantified using Salmon [v.X.X.X] (Patro et al. 2017). To filter out low-confidence hits and potential noise, species presence was scored only when for genes that showed transcriptional activity, defined as a minimum threshold of 10 mapped reads per sample per reference species.

### Expression of mycorrhizal and drought-related gene markers in argan

We identified gene markers related to drought and the mycorrhizal symbiosis on our argan transcriptome. We used a list of core mycorrhizal genes as a set of mycorrhizal gene-markers. We then used a blast method to identify homologs between the mycorrhizal gene-markers and argan. For drought-related gene markers, we used a list of genes related to drought from a comparative analysis of *Arabidopsis thaliana* and the xerophytic species *Reaumuria soongorica* (Shi et al. 2013). We used OMA 2.6.1 (Altenhoff et al. 2019) to identify orthologs between *Arabidopsis thaliana* and argan transcriptomes. We then identified the list of drought-related genes in arabidopsis and the corresponding orthologs in argan.

## Acknowledgments

We are grateful to BGI Tech Solutions for performing the RNA sequencing and to the Bern High-Performance Computing (HPC) facility for providing the computational resources required for bioinformatic analysis. We also thank Axelle Raisin and Nathalie Judkins for their valuable comments and critical reading of the manuscript.

## Author Contributions

AbdE and AurE performed experiments, LF and IM carried out bioinformatics and statistics analysis, AbdE, AQ, IM, and DR conceived the project; all authors contributed to the manuscript.

## Data availability statement

All data are available in the Supplementary Figures and Tables. All raw sequence data were submitted to the EBI/ENA repository (accession Project No.: PRJEB60382).

## Supplementary Figures

**Figure S1. Sampling of native mycorrhizal inocula in the Arganeraie.**

At each sampling site, three soil and root samples were collected and pooled from individual argan trees. Indicated are the exact locations of sampling sites S1-S10 with detailed coordinates and altitude (meters above sea level).

**Figure S2. Mycorrhizal colonization with native inocula under drought.**

Mycorrhizal colonization was determined in plants grown under controlled conditions with native Moroccan inocula S1-S10 (see **Figure S1**). For comparison, inocula from *R. irregularis* DAOM197198 (Ri) and the drought-adapted AM fungus *D. omaniana* (Do) were included, as well as non-mycorrhizal controls (NM). Plants were cultured for 8 weeks after inoculation under well-watered conditions (blue), or drought (red columns). Mean values (n=5) ± SE are shown. Significant effects (*P* ≤ 0.05 one-way ANOVA and Tukey’s test) of drought on root colonization are indicated * *P* < 0.05; ** *P* < 0.01; *** *P* < 0.001.

**Figure S3. Global growth parameters of plants colonized by native Moroccan AMF inocula.**

Shoot height **(a)**, shoot dry weight **(b)**, root dry weight **(c)** and leaf area **(d)** of plants inoculated with 10 Moroccan inocula (S1-S10), and with the reference AMF species *R. irregularis* (RI) and *D. omaniana* (DO) and grown under well-watered or drought conditions, respectively. Horizontal lines indicate the level of non-mycorrhizal plants (NM) under drought as reference for the protective effect of the various mycorrhizal inocula under drought. Mean values (n=5) ± SE are shown. Significant effects of drought (*P* ≤ 0.05; one-way ANOVA and Tukey’s test) on the respective traits are indicated with black asterisks (* *P* < 0.05; ** *P* < 0.01; *** *P* < 0.001). Colored asterisks indicate significant differences of the inoculated plants vs. non-mycorrhizal controls under well-watered conditions (green asterisks), and under drought (red asterisks), respectively (* *P* < 0.05; ** *P* < 0.01; *** *P* < 0.001).

**Figure S4. Growth dynamics of argan colonized by *R. irregularis* under drought.**

Shoot height of plants inoculated with *R. irregularis* (Ri), with native inoculum S10 (Ni), or non-mycorrhizal (NM) under well-watered or drought conditions, respectively. Mean values (n=5) ± SE are shown.

**Figure S5. Effects of mycorrhizal colonization on water relations.**

Accumulated water uptake **(a)**, leaf relative water content (RWC) **(b)**, and stomatal conductance **(c)** of plants inoculated with 10 Moroccan inocula (S1-S10), and with the reference AMF species *R. irregularis* (RI) and *D. omaniana* (DO) and grown under well-watered (blue columns) or drought conditions (red columns). Horizontal lines indicate the level of non-mycorrhizal plants (NM) under drought as reference for the protective effect of the various mycorrhizal inocula under drought. Mean values (n=5) ± SE are shown. Significant effects of drought (*P* ≤ 0.05; one-way ANOVA and Tukey’s test) on the respective traits are indicated with black asterisks (* *P* < 0.05; ** *P* < 0.01; *** *P* < 0.001). Colored asterisks indicate significant differences of the inoculated plants vs. non-mycorrhizal controls under well-watered conditions (green asterisks), and under drought (red asterisks), respectively (* *P* < 0.05; ** *P* < 0.01; *** *P* < 0.001).

**Figure S6. Principle component analysis of phenotypic traits of plants with or without *R. irregularis* and under well-watered or drought conditions.**

Notable most effects were observed in the horizontal dimension (Dim1), which determined 52% of the effects. Mycorrhizal status caused a shift to the upper right, while drought caused an opposite shift to the left. Annotated arrows indicate in which direction the cloud as shifted by the respective factor.

**Figure S7. Mycorrhizal effects on various growth-related parameters in well-watered plants inoculated with three native Moroccan inocula (S3, S8, S10)**

Relative difference (%) in 17 growth-related parameters determined in mycorrhizal plants vs. non-mycorrhizal controls grown under well-watered conditions. Eight weeks after inoculation with native inocula S3, S8 or S10, the following parameters were determined: root dry weight, shoot dry weight, shoot height, number of lateral branches (SN), leaf area, relative water content of the leaves (RWC), accumulated water uptake during the experiment, the content of proline, soluble sugars, and chlorophyll (Chlo) in the leaves, non-photochemical quenching (PhiNPQ), quantum yield of photosystem II (Phi2), the levels of malondialdehyde (MDA) and hydrogen peroxide (H2O2, as well as the activities of (POD), and catalase (CAT). Bars represent mean values (n=5); significance levels (one-way ANOVA and Tukey’s test) are indicated as follows * *P* < 0.05; ** *P* < 0.01.

**Figure S8. Mycorrhizal effects on various growth-related parameters of plants inoculated with three native Moroccan inocula (S3, S8, S10) and grown under drought conditions.**

Relative difference (%) in 17 growth-related parameters determined in mycorrhizal plants vs. non-mycorrhizal controls grown under drought. Eight weeks after inoculation with native inocula S3, S8 or S10, the following parameters were determined: root dry weight, shoot dry weight, shoot height, number of lateral branches (SN), leaf area, relative water content of the leaves (RWC), accumulated water uptake during the experiment, the content of proline, soluble sugars, and chlorophyll (Chlo) in the leaves, non-photochemical quenching (PhiNPQ), quantum yield of photosystem II (Phi2), the levels of malondialdehyde (MDA) and hydrogen peroxide (H2O2, as well as the activities of (POD), and catalase (CAT). Bars represent mean values (n=5); significance levels (one-way ANOVA and Tukey’s test) are indicated as follows * *P* < 0.05; ** *P* < 0.01.

**Figure S9. Global clustering analysis of plants grown under the two water regimes with four different inocula.**

Heatmap shows fold change (log2) of the 17 growth- and stress-related parameters relative to the respective non-mycorrhizal controls. Note clustering according to water conditions, and the general reduction by drought of parameters related to growth, and a general induction of parameters related to stress.

**Figure S10. Histochemical DAB staining of H_2_O_2_ in the leaves.**

Staining of leaves with diaminobenzidine (DAB) for H_2_O_2_ detection in argan plants grown under well-watered conditions and under drought, growing without AMF inoculum (NM), with three native inocula (S3, S8 and S10), or with *R. irregularis* (RI). No DAB staining was observed except in the positive control wounded by forceps.

**Figure S11. Drought stress in maize.**

**(a-c)** Shoot appearance and fresh weight of maize plants grown under well-watered conditions **(a)**, and drought **(b)**, and respective shoot fresh weight **(c)**.

**(d-g)** Histochemical DAB staining of H_2_O_2_ in the leaves of maize plants grown under well-watered conditions **(d,e)** or drought **(f,g)**.

**Figure S12. Principle-Component analysis of RNAseq data.**

Plants wer grown with or without mycorrhizal inoculum (AMF, noAMF, respectively), ander well-watered conditions (80% FC), or under drought (20% FC). Mycorrhizal samples appeared well clustered, while non-mycorrhizal samples were rather spread.

**Figure S13. AM fungal composition of inoculum S8.**

Fungal RNA-Seq reads were assigned to 24 AM fungal species for which a genome sequence was available. The heatmap indicates relative frequency. Fungal reads were restricted to inoculated plants (AM). Inoculum S8 contained mainly reads assigned to the genera Rhizophagus, Funnelliformis, Glomus, Claroideoglomus and Paraglomus.

**Figure S14. Regulation of drought-responsive genes in non-mycorrhizal controls.** Clustering of the top 0.5% genes based on the p-values for significance of induction. Shown are genes induced or repressed by drought in non-mycorrhizal samples. The heatmap shows relative expression level.

**Figure S15. Regulation of drought-responsive genes in mycorrhizal roots.**

Clustering of the top 0.5% genes based on the p-values for significance of induction. Shown are genes induced or repressed by drought in mycorrhizal samples. The heatmap shows relative expression level.

**Figure S16. Regulation of AM-responsive genes under well-watered conditions.** Clustering of the top 0.5% genes based on the p-values for significance of induction. Shown are genes induced or repressed in mycorrhizal roots under well-watered conditions. The heatmap shows relative expression level.

**Figure S17. Regulation of AM-responsive genes under drought.**

Clustering of the top 0.5% genes based on the p-values for significance of induction. Shown are genes induced or repressed in mycorrhizal roots under drought. The heatmap shows relative expression level.

**Figure S18. Semantic space visualization of Gene Ontology (GO) term enrichment.** Scatter plots summarizing the enriched GO biological processes associated with the drought effect **(a)** and the mycorrhizal effect **(b)**, generated with GO-Figure. Enriched GO terms are projected into a two-dimensional semantic space (Semantic dimensions X and Y) based on the semantic similarity of their GO categories. Spatially adjacent points represent biological processes that are functionally closely related. The color of each bubble corresponds to the statistical significance of the enrichment. The size of the bubbles represent the size of each enriched GO-term.

**Table S1. Phenotypic characterization of argan.**

Growth, physiological and mycorrhizal colonization variables measured on argan in drought and mycorrhizal colonization conditions.

**Table S2. Global statistics of RNA sequencing**

RNAseq raw reads, filtered reads and number of genes detected.

**Table S3. Genes affected by drought and mycorrhizal status.**

Complete results from the RNAseq experiment with the annotated argan transcriptome. Functional annotation of genes was produced by eggNOG and the closest homologs of the argan genes in the model species *Medicago truncatula*, *Oryza sativa* (rice), and *Arabidopsis thaliana* are provided. Lists of expression ratios are organized in 4 tables according to the different comparisons performed in the RNAseq analysis. 1) Drought effect in mycorrhizal roots, 2) Drought effect whitout mycorrhiza, 3) Mycorrhizal effect in drought conditions and 4) Mycorrhizal effect in well-watered conditions.

**Table S4. Argan homologues of AM-related genes**

The five to ten closest argan homologs of known mycorrhizal marker genes were identified by protein Blast using mycorrhiza-related proteins described by Rich et al., 2017. Asterisks indicate PT4 homologs that were detected in haplotype 2, but not in haplotype 1 of argan.

**Table S5. Argan orthologs of Arabidopsis drought-related genes**

Argan orthologs of Arabidopsis genes assigned to stress avoidance and stress tolerance, respectively, in the response to drought as defined by Shi et al. (2013)

## References

Aebi H (1984) Catalase *in vitro*. Methods Enzymol 105:121–126

Alexa A, Rahnenführer J, Lengauer T (2006) Improved scoring of functional groups from gene expression data by decorrelating GO graph structure. Bioinformatics 22 (13):1600–1607. doi:10.1093/bioinformatics/btl140

Altenhoff AM, Levy J, Zarowiecki M, Tomiczek B, Vesztrocy AW, Dalquen DA, Müller S, Telford MJ, Glover NM, Dylus D, Dessimoz C (2019) OMA standalone: Orthology inference among public and custom genomes and transcriptomes. Genome Research 29 (7):1152–1163. 10.1101/gr.243212.118

Arnon DI (1949) Copper enzymes in isolated chloroplasts - polyphenoloxidase in *Beta vulgaris*. Plant Physiol 24 (1):1–15. doi:10.1104/pp.24.1.1

Augé RM, Toler HD, Saxton AM (2015) Arbuscular mycorrhizal symbiosis alters stomatal conductance of host plants more under drought than under amply watered conditions: a meta-analysis. Mycorrhiza 25 (1):13–24. doi:10.1007/s00572-014-0585-4

Bakhat N, Vielba-Fernández A, Padilla-Roji I, Martínez-Cruz J, Polonio A, Fernández-Ortuño D, Pérez-García A (2023) Suppression of chitin-triggered immunity by plant fungal pathogens: A case study of the cucurbit powdery mildew fungus *Podosphaera xanthii*. Journal of Fungi 9 (7). doi:10.3390/jof9070771

Bandurska H (2022) Drought stress responses: Coping strategy and resistance. Plants 11 (7). doi:10.3390/plants11070922

Barradas MCD, Zunzunegui M, Esquivias MP, Boutaleb S, Valera-Burgos J, Tagma T, Ain-Lhout F (2013) Some secrets of *Argania spinosa* water economy in a semiarid climate. Natural Product Communications 8 (1):11–14

Bates LS, Waldren RP, Teare ID (1973) Rapid determination of free proline for water-stress studies. Plant Soil 39 (1):205–207. doi:10.1007/bf00018060

Benhiba L, Fouad MO, Essahibi A, Ghoulam C, Qaddoury A (2015) Arbuscular mycorrhizal symbiosis enhanced growth and antioxidant metabolism in date palm subjected to long-term drought. Trees-Structure and Function 29 (6):1725–1733. doi:10.1007/s00468-015-1253-9

Berruti A, Lumini E, Balestrini R, Bianciotto V (2016) Arbuscular mycorrhizal fungi as natural biofertilizers: Let’s benefit from past successes. Frontiers in Microbiology 6. doi:10.3389/fmicb.2015.01559

Bravo A, York T, Pumplin N, Mueller LA, Harrison MJ (2016) Genes conserved for arbuscular mycorrhizal symbiosis identified through phylogenomics. Nature Plants 2 (2):15208. doi:10.1038/nplants.2015.208

Breuillin F, Schramm J, Hajirezaei M, Ahkami A, Favre P, Druege U, Hause B, M. B, Kretzschmar T, Bossolini E, Kuhlemeier C, Martinoia E, Franken P, Scholz U, Reinhardt D (2010) Phosphate systemically inhibits development of arbuscular mycorrhiza in *Petunia hybrida* and represses genes involved in mycorrhizal functioning. Plant J 64 (6):1002–1017

Brunner I, Herzog C, Dawes MA, Arend M, Sperisen C (2015) How tree roots respond to drought. Frontiers in Plant Science 6. doi:10.3389/fpls.2015.00547

Cantalapiedra CP, Hernández-Plaza A, Letunic I, Bork P, Huerta-Cepas J (2021) eggNOG-mapper v2: Functional Annotation, Orthology Assignments, and Domain Prediction at the Metagenomic Scale. Molecular Biology and Evolution 38 (12):5825–5829. doi:10.1093/molbev/msab293

Chakhchar A, Ben Salah I, El Kharrassi Y, Filali-Maltouf A, El Modafar C, Lamaoui M (2022) Agro-fruit-forest systems based on argan tree in Morocco: A review of recent results. Frontiers in Plant Science 12. doi:10.3389/fpls.2021.783615

Chakhchar A, Haworth M, El Modafar C, Lauteri M, Mattioni C, Wahbi S, Centritto M (2017) An assessment of genetic diversity and drought tolerance in argan tree (*Argania spinosa*) populations: Potential for the development of improved drought tolerance. Frontiers in Plant Science 8:1–11. 10.3389/fpls.2017.00276

Chakhchar A, Lamaoui M, Aissam S, Ferradous A, Wahbi S, El Mousadik A, Ibnsouda-Koraichi S, Filali-Maltouf A, El Modafar C (2018) Physiological and biochemical mechanisms of drought stress tolerance in the argan tree. In: Ahmad P, Ahanger MA, Singh VP, Tripathi DK, Alam P, Alyemeni MN (eds) Plant Metabolites and Regulation Under Environmental Stress. Academic Press, pp 311–322. doi:10.1016/B978-0-12-812689-9.00015-7

Chakhchar A, Soufiani M, Lamaoui M, Ferradous A, Wahbi S, El Mousadik A, Ibnsouda-Koraichi S, Filali-Maltouf A, El Modafar C (2024) Delineating drought-induced antioxidative traits as potential mechanisms for climate resilience in *Argania spinosa*. Notulae Scientia Biologogiae 16 (3):12000. doi:10.15835/nsb16312000

Cheeseman JM (2006) Hydrogen peroxide concentrations in leaves under natural conditions. J Exp Bot 57 (10):2435–2444. doi:10.1093/jxb/erl004

Chen M, Arato M, Borghi L, Nouri E, Reinhardt D (2018a) Beneficial services of arbuscular mycorrhizal fungi - From ecology to application. Frontiers in Plant Science 9:1270. doi:10.3389/fpls.2018.01270

Chen SF, Zhou YQ, Chen YR, Gu J (2018b) fastp: an ultra-fast all-in-one FASTQ preprocessor. Bioinformatics 34 (17):884–890. doi:10.1093/bioinformatics/bty560

Chen X, Zhao CC, Yun P, Yu M, Zhou MX, Chen ZH, Shabala S (2024) Climate-resilient crops: Lessons from xerophytes. Plant J 117 (6):1815–1835. doi:10.1111/tpj.16549

Chieb M, Gachomo EW (2023) The role of plant growth promoting rhizobacteria in plant drought stress responses. Bmc Plant Biology 23 (1). doi:10.1186/s12870-023-04403-8

Cruz de Carvalho MH (2008) Drought stress and reactive oxygen species. Plant Signaling & Behavior 3:156–165

Daudi A, O’Brien JA (2012) Detection of hydrogen peroxide by DAB staining in Arabidopsis leaves. BioProtocol 2:1–4

Díaz-Barradas MC, Zunzunegui M, Ain-Lhout F, Jáuregui J, Boutaleb S, Alvarez-Cansino L, Esquivias MP (2010) Seasonal physiological responses of *Argania spinosa* tree from mediterranean to semi-arid climate. Plant Soil 337 (1-2):217–231. doi:10.1007/s11104-010-0518-8

Dobin A, Davis CA, Schlesinger F, Drenkow J, Zaleski C, Jha S, Batut P, Chaisson M, Gingeras TR (2013) STAR: ultrafast universal RNA-seq aligner. Bioinformatics 29 (1):15–21. doi:10.1093/bioinformatics/bts635

El Idrissi H, Gkanogiannis A, Iraqi D, Khoulassa S, Fokar M, Badaoui B, Moussadek R, Mentag R, Khayi S (2026) A phased, near-telomere-to-telomere chromosome-scale reference genome of the Moroccan argan tree. Scientific Data 13 (1). doi:10.1038/s41597-026-06615-7

Essahibi A, Benhiba L, Babram MA, Ghoulam C, Qaddoury A (2018) Influence of arbuscular mycorrhizal fungi on the functional mechanisms associated with drought tolerance in carob (Ceratonia siliqua L.). Trees-Structure and Function 32 (1):87–97. doi:10.1007/s00468-017-1613-8

Essahibi A, Benhiba L, Fouad MO, Babram MA, Ghoulam C, Qaddoury A (2019) Responsiveness of carob (*Ceratonia siliqua* L.) plants to arbuscular mycorrhizal symbiosis under different phosphate fertilization levels. J Plant Growth Regul 38 (4):1243–1254. doi:10.1007/s00344-019-09929-6

Etesami H, Otabek U, Zahro B, Laziz Y, Sevara N (2026) Arbuscular mycorrhizal fungi in desert ecosystems: Adaptive mechanisms to co-occurring drought, temperature, and salinity stress. Fungal Biology Reviews 55. doi:10.1016/j.fbr.2026.100472

Evelin H, Devi TS, Gupta S, Kapoor R (2019) Mitigation of salinity stress in plants by arbuscular mycorrhizal symbiosis: Current understanding and new challenges. Frontiers in Plant Science 10. doi:10.3389/fpls.2019.00470

Favre P, Bapaume L, Bossolini E, Delorenzi L, Falquet L, Reinhardt D (2014) A novel bioinformatics pipeline to discover genes related to arbuscular mycorrhizal symbiosis based on their evolutionary conservation pattern among higher plants. BMC Plant Biology 14:333

Formenti C, Mauromicale G, Pandino G, Lombardo S (2026) Arbuscular mycorrhizal fungi mitigate crop multi-stresses under mediterranean climate: A systematic review. Agronomy-Basel 16 (1). doi:10.3390/agronomy16010113

Fuchslueger L, Bahn M, Fritz K, Hasibeder R, Richter A (2014) Experimental drought reduces the transfer of recently fixed plant carbon to soil microbes and alters the bacterial community composition in a mountain meadow. New Phytol 201 (3):916–927. doi:10.1111/nph.12569

Ganoudi M, El Malahi S, Manan N, Ibriz M, Calonne-Salmon M, Declerck S (2025) Drought resistance of *Argania spinosa* L. colonized by the arbuscular mycorrhizal fungus *Rhizophagus irregularis* varies according to accession. Frontiers in Plant Science 16. doi:10.3389/fpls.2025.1678553

Gobbato E, Wang E, Higgins G, Bano SA, Henry C, Schultze M, Oldroyd GEd (2013) RAM1 and RAM2 function and expression during arbuscular mycorrhizal symbiosis and *Aphanomyces euteiches* colonization. Plant Signaling and Behavior 8 (10):e26049

Hernández JA, Almansa MS (2002) Short-term effects of salt stress on antioxidant systems and leaf water relations of pea leaves. Physiol Plant 115 (2):251–257. doi:10.1034/j.1399-3054.2002.1150211.x

Heyduk K (2022) Evolution of Crassulacean acid metabolism in response to the environment: past, present, and future. Plant Physiol 190 (1):19–30. doi:10.1093/plphys/kiac303

Hori K, Wada A, Shibuta T (1997) Changes in phenoloxidase activities of the galls on leaves of Ulmus davidana formed by *Tetraneura fusiformis* (Homoptera: Eriosomatidae). Applied Entomology and Zoology 32 (2):365–371. doi:10.1303/aez.32.365

Howard CC, Folk RA, Beaulieu JM, Cellinese N (2019) The monocotyledonous underground: global climatic and phylogenetic patterns of geophyte diversity. Am J Bot 106 (6):850–863. doi:10.1002/ajb2.1289

Irigoyen JJ, Emerich DW, Sanchezdiaz M (1992) Water-stress induced changes in concentrations of proline and total soluble sugars in nodulated alfalfa (*Medicago sativa*) plants. Physiol Plant 84 (1):55–60. doi:10.1034/j.1399-3054.1992.840109.x

Kirchhoff M, Engelmann L, Zimmermann LL, Seeger M, Marzolff I, Hssaine AA, Ries JB (2019) Geomorphodynamics in argan woodlands, south Morocco. Water 11 (10). doi:10.3390/w11102193

Klironomos JN (2003) Variation in plant response to native and exotic arbuscular mycorrhizal fungi. Ecology 84 (9):2292–2301

Levitt J (1972) Responses of Plants to Environmental Stresses. Academic Press, San Diego, CA, USA

Li XM, Xi BY, Wu XC, Choat B, Feng JC, Jiang MK, Tissue D (2022) Unlocking drought-induced tree mortality: Physiological mechanisms to modeling. Frontiers in Plant Science 13. doi:10.3389/fpls.2022.835921

Liao HX, Zhao HJ, Chen ZH, Peng SL (2025) Drought response of trees: differences across mycorrhizal type at the global scale. Oikos 2025 (10). doi:10.1002/oik.11257

Liao Y, Smyth GK, Shi W (2014) featureCounts: an efficient general purpose program for assigning sequence reads to genomic features. Bioinformatics 30 (7):923–930. doi:10.1093/bioinformatics/btt656

Lionello P, Scarascia L (2018) The relation between climate change in the Mediterranean region and global warming. Regional Environmental Change 18 (5):1481–1493. doi:10.1007/s10113-018-1290-1

Liu X, Quan WL, Bartels D (2022) Stress memory responses and seed priming correlate with drought tolerance in plants: an overview. Planta 255 (2). doi:10.1007/s00425-022-03828-z

Love MI, Huber W, Anders S (2014) Moderated estimation of fold change and dispersion for RNA-seq data with DESeq2. Genome Biology 15 (12). doi:10.1186/s13059-014-0550-8

Lugtenberg B, Kamilova F (2009) Plant-growth-promoting rhizobacteria. Annual Review of Microbiology 63:541–556. doi:10.1146/annurev.micro.62.081307.162918

Lybbert TJ, Aboudrare A, Chaloud D, Magnan N, Nash M (2011) Booming markets for Moroccan argan oil appear to benefit some rural households while threatening the endemic argan forest. Proc Natl Acad Sci U S A 108 (34):13963–13968. 10.1073/pnas.1106382108

Madouh TA, Suleiman MK, Quoreshi AM, Davidson MK (2025) Diversity of arbuscular mycorrhiza fungi in the arid desert ecosystems of Kuwait: Detection and identification from perennial native grass roots. Diversity-Basel 17 (2). doi:10.3390/d17020130

Manley BF, Lotharukpong JS, Barrera-Redondo J, Llewellyn T, Yildirir G, Sperschneider J, Corradi N, Paszkowski U, Miska EA, Dallaire A (2023) A highly contiguous genome assembly reveals sources of genomic novelty in the symbiotic fungus *Rhizophagus irregularis*. G3-Genes Genomes Genetics 13 (6). doi:10.1093/g3journal/jkad077

Marks RA, Farrant JM, McLetchie DN, VanBuren R (2021) Unexplored dimensions of variability in vegetative desiccation tolerance. Am J Bot 108 (2):346–358. doi:10.1002/ajb2.1588

Mateus ID, Essahibi A, Nicholson P, Hijri M, Qaddoury A, Falquet L, Reinhardt D (2025) Chromosome-level phased genome assembly of the argan tree *Sideroxylon spinosum*. Scientific Data 12:1430

Meddich A, Oihabi A, Abbas Y, Bizid E (2000) Rôle des champignons mycorhiziens à arbuscules de zones arides dans la résistance du trèfle (*Trifolium alexandrinum* L.) au déficit hydrique. Agronomie 20:283–295. doi:10.1051/agro:2000127

Metze D, Schnecker J, Canarini A, Fuchslueger L, Koch BJ, Stone BW, Hungate BA, Hausmann B, Schmidt H, Schaumberger A, Bahn M, Kaiser C, Richter A (2023) Microbial growth under drought is confined to distinct taxa and modified by potential future climate conditions. Nature Communications 14 (1). doi:10.1038/s41467-023-41524-y

Montoliu-Nerin M, Sanchez-Garcia M, Bergin C, Kutschera VE, Johannesson H, Bever JD, Rosling A (2021) In-depth phylogenomic analysis of arbuscular mycorrhizal fungi based on a comprehensive set of *de novo* genome assemblies. Front Fungal Biol 2:716385. doi:10.3389/ffunb.2021.716385

Moukrim S, Lahssini S, Rhazi M, Alaoui HM, Benabou A, Wahby I, El Madihi M, Arahou M, Rhazi L (2019) Climate change impacts on potential distribution of multipurpose agro-forestry species: *Argania spinosa* (L.) Skeels as case study. Agroforestry Systems 93 (4):1209–1219. doi:10.1007/s10457-018-0232-8

Msanda F, El Aboudi A, Peltier J (2005) Biodiversity and biogeography of Moroccan argan tree communities. Cahiers Agricultures 14:357–364

Nouaim R, Chaussod R (1994) Mycorrhizal dependency of micropropagated argan tree (*Argania-spinosa*). 1. Growth and biomass production. Agroforestry Systems 27 (1):53–65. doi:10.1007/bf00704834

Nour MM, Aljabi HR, Al-Huqail AA, Horneburg B, Mohammed AE, Alotaibi MO (2024) Drought responses and adaptation in plants differing in life-form. Frontiers in Ecology and Evolution 12. doi:10.3389/fevo.2024.1452427

Ostrowska A, Hura K, Hura T (2024) Accumulation of hydrogen peroxide in flag leaves induces effective regeneration of triticale during rehydration after water stress. J Plant Growth Regul 43 (10):3560–3569. doi:10.1007/s00344-024-11333-8

Park EJ, Kim TH (2021) Thaumatin-like genes function in the control of both biotic stress signaling and ABA signaling pathways. Biochem Biophys Res Commun 567:17–21. doi:10.1016/j.bbrc.2021.06.012

Patro R, Duggal G, Love MI, Irizarry RA, Kingsford C (2017) Salmon provides fast and bias-aware quantification of transcript expression. Nature Methods 14 (4):417-+. doi:10.1038/nmeth.4197

Phillips JM, Hayman DS (1970) Improved procedures for clearing roots and staining parasitic and vesicular-arbuscular mycorrhizal fungi for rapid assessment of infection. Transactions of the British Mycological Society 55:158–161

Pieterse CMJ (2025) The extended plant immune system. Mol Plant-Microbe Interact 38 (6):780–795. doi:10.1094/mpmi-10-25-0144-hh

Pimprikar P, Carbonnel S, Paries M, Katzer K, Klingl V, Bohmer MJ, Karl L, Floss DS, Harrison MJ, Parniske M, Gutjahr C (2016) A CCaMK-CYCLOPS-DELLA complex activates transcription of RAM1 to regulate arbuscule branching. Current Biology 26 (8):987–998

Reijnders MJ, Waterhouse RM (2021) Summary visualizations of gene ontology terms with GO-Figure!. Front Bioinform 1:638255. doi:10.3389/fbinf.2021.638255

Rich MK, Courty P-E, Roux C, Reinhardt D (2017) Role of the GRAS transcription factor ATA/RAM1 in the transcriptional reprogramming of arbuscular mycorrhiza in *Petunia hybrida*. BMC Genomics 18:589

Rupp O, Roessner C, Lederer-Ponzer N, Wollenweber TE, Becker A, Lamaoui M (2024) Genome of *Argania spinosa* L.: insights into oil production and the tocopherol biosynthesis pathway. Genetic Resources and Crop Evolution 71 (8):4027–4042. 10.1007/s10722-024-01931-6

Saeed Ur R, Khalid M, Hui N, Kayani SI, Tang KX (2020) Diversity and versatile functions of metallothioneins produced by plants: A review. Pedosphere 30 (5):577–588. doi:10.1016/s1002-0160(20)60022-4

Salminen TA, Blomqvist K, Edqvist J (2016) Lipid transfer proteins: classification, nomenclature, structure, and function. Planta 244 (5):971–997. doi:10.1007/s00425-016-2585-4

Savary R, Masclaux FG, Wyss T, Droh G, Corella JC, Machado AP, Morton JB, Sanders IR (2018) A population genomics approach shows widespread geographical distribution of cryptic genomic forms of the symbiotic fungus *Rhizophagus irregularis*. ISME Journal 12 (1):17–30. doi:10.1038/ismej.2017.153

Schimel JP (2018) Life in dry soils: Effects of drought on soil microbial communities and processes. In: Futuyma DJ (ed) Annual Review of Ecology, Evolution, and Systematics, vol 49. Annual Review of Ecology Evolution and Systematics. pp 409–432. doi:10.1146/annurev-ecolsys-110617-062614

Shi Y, Yan X, Zhao PS, Yin HX, Zhao X, Xiao HL, Li XR, Chen GX, Ma XF (2013) Transcriptomic analysis of a tertiary relict plant, extreme xerophyte *Reaumuria soongorica* to identify genes related to drought adaptation. Plos One 8 (5). doi:10.1371/journal.pone.0063993

Smith SE, Read DJ (2008) Mycorrhizal Symbiosis. 3rd edition edn. Academic Press, New York

Symanczik S, Blaszkowski J, Chwat G, Boller T, Wiemken A, Al-Yahya’ei MN (2014) Three new species of arbuscular mycorrhizal fungi discovered at one location in a desert of Oman: *Diversispora omaniana*, *Septoglomus nakheelum* and *Rhizophagus arabicus*. Mycologia 106 (2):243–259. doi:10.3852/106.2.243

Talaat NB, Shawky BT (2014) Protective effects of arbuscular mycorrhizal fungi on wheat (*Triticum aestivum* L.) plants exposed to salinity. Environ Exp Bot 98:20–31. doi:10.1016/j.envexpbot.2013.10.005

Tariq A, Gao YJ, Zeng FJ, Sardans J, Ahmed Z, Graciano C, Hughes AC, Peñuelas J (2026) Guardians of arid lands: deep-rooted defense against desertification and climate change. Trends Plant Sci 31 (3):295–312. doi:10.1016/j.tplants.2025.10.009

Team Rc R: A Language and Environment for Statistical Computing.

Terral JF, Bédé G, Paradis L, Ivorra S, Ros J, Girard V, Ater M, Limier B, Kassout J (2025) Understanding how the Argan tree adapts its performance in a heterogeneous environment using wood anatomical and leaf functional traits. J Arid Environ 231. doi:10.1016/j.jaridenv.2025.105473

Timzioura R, Ezzine S, Benomar L, Lamhamedi MS, Ettaqy A, El Abidine AZ, Zaher H, Khasa DP, Pepin S, Abbas Y (2025) Bibliometric Analysis of Argan (Argania spinosa (L.) Skeels) Research: Scientific Trends and Strategic Directions for Climate-Resilient Ecosystem Management. Forests 16 (6). doi:10.3390/f16060892

Tisserant E, Malbreil M, Kuo A, Kohler A, Symeonidi A, Balestrini R, Charron P, Duensing N, Frey NFD, Gianinazzi-Pearson V, Gilbert LB, Handa Y, Herr JR, Hijri M, Koul R, Kawaguchi M, Krajinski F, Lammers PJ, Masclauxm FG, Murat C, Morin E, Ndikumana S, Pagni M, Petitpierre D, Requena N, Rosikiewicz P, Riley R, Saito K, Clemente HS, Shapiro H, Van Tuinen D, Bécard G, Bonfante P, Paszkowski U, Shachar-Hill YY, Tuskan GA, Young PW, Sanders IR, Henrissat B, Rensing SA, Grigoriev IV, Corradi N, Roux C, Martin F (2013) Genome of an arbuscular mycorrhizal fungus provides insight into the oldest plant symbiosis. Proc Natl Acad Sci U S A 110 (50):20117–20122. doi:10.1073/pnas.1313452110

Trouvelot A, Kough JL, Gianinazzi-Pearson V (1986) Mesure du taux de mycorhization VA d’un système radiculaire. Recherche de méthodes d’éstimation ayant une signification fonctionelle.. In: Les mycorhizes: physiologie et génétique. ESM/SEM, Dijon, 1-5 July. INRA Press, Paris, pp 217–222

Usman MG, Rafii MY, Martini MY, Yusuff OA, Ismail MR, Miah G (2017) Molecular analysis of Hsp70 mechanisms in plants and their function in response to stress. Biotechnology and Genetic Engineering Reviews 33 (1):26–39. doi:10.1080/02648725.2017.1340546

Vasar M, Davison J, Sepp SK, Öpik M, Moora M, Koorem K, Meng YM, Oja J, Akhmetzhanova AA, Al-Quraishy S, Onipchenko VG, Cantero JJ, Glassman SI, Hozzein WN, Zobel M (2021) Arbuscular mycorrhizal fungal communities in the soils of desert habitats. Microorganisms 9 (2). doi:10.3390/microorganisms9020229

Velikova V, Yordanov I, Edreva A (2000) Oxidative stress and some antioxidant systems in acid rain-treated bean plants - Protective role of exogenous polyamines. Plant Sci 151 (1):59–66. doi:10.1016/s0168-9452(99)00197-1

Yao LY, Cheng X, Gu ZY, Huang W, Li S, Wang LB, Wang YF, Xu P, Ma H, Ge XC (2018) The AWPM-19 family protein OsPM1 mediates abscisic acid influx and drought response in rice. Plant Cell 30 (6):1258–1276. doi:10.1105/tpc.17.00770

