## Supplemental Figures for "Arbuscular mycorrhizal symbiosis increases drought resistance in the xerophytic argan tree (*Sideroxylon spinosum*)"

### Supplementary Figures

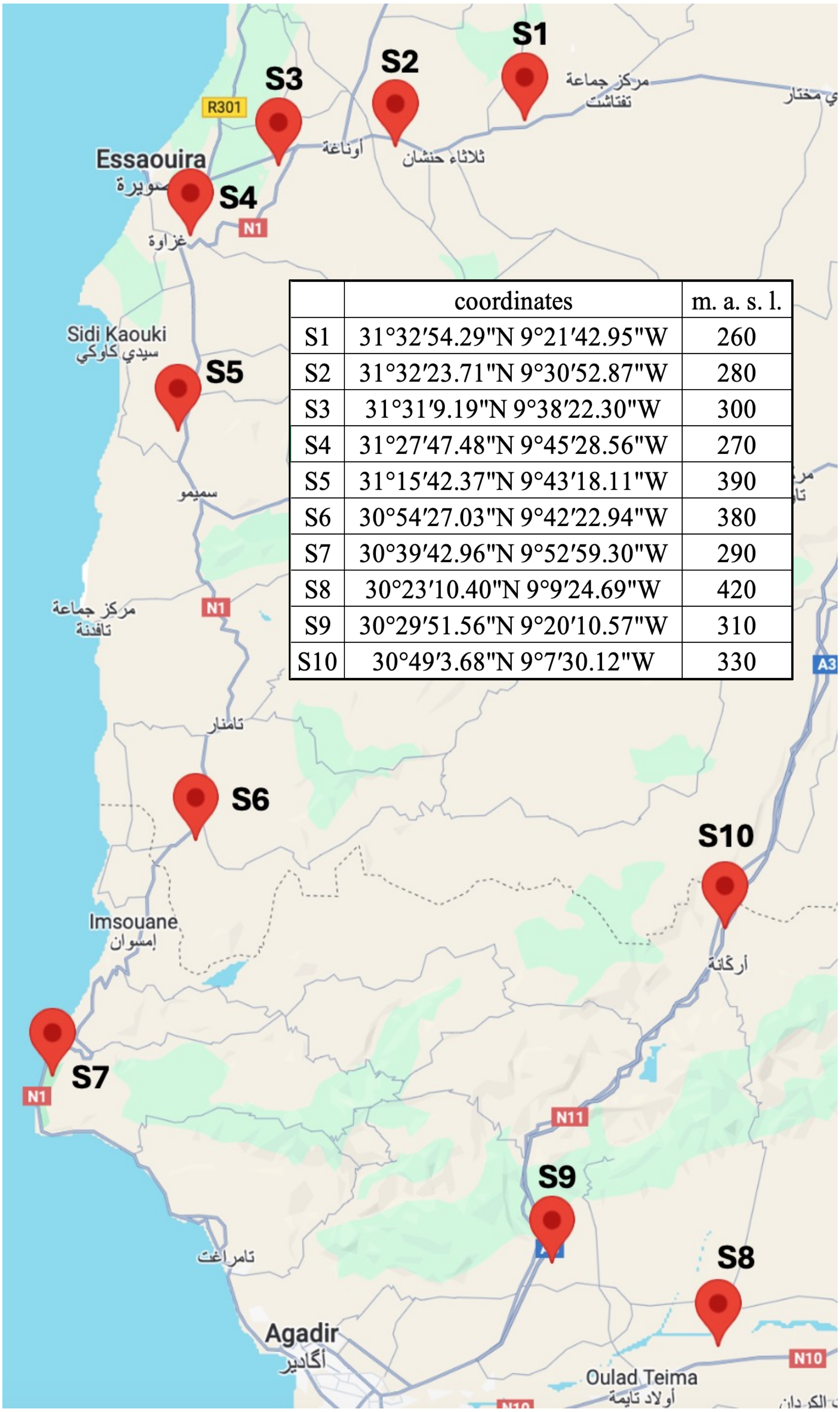


| | coordinates | m. a. s. l. |
|---|---|---|
| S1 | 31°32′54.29"N 9°21′42.95"W | 260 |
| S2 | 31°32′23.71"N 9°30′52.87"W | 280 |
| S3 | 31°31′9.19"N 9°38′22.30"W | 300 |
| S4 | 31°27′47.48"N 9°45′28.56"W | 270 |
| S5 | 31°15′42.37"N 9°43′18.11"W | 390 |
| S6 | 30°54′27.03"N 9°42′22.94"W | 380 |
| S7 | 30°39′42.96"N 9°52′59.30"W | 290 |
| S8 | 30°23′10.40"N 9°9′24.69"W | 420 |
| S9 | 30°29′51.56"N 9°20′10.57"W | 310 |
| S10 | 30°49′3.68"N 9°7′30.12"W | 330 |

**Figure S1. Sampling of native mycorrhizal inocula in the Arganeraie.**
At each sampling site, three soil and root samples were collected and pooled from individual argan trees. Indicated are the exact locations of sampling sites S1-S10 with detailed coordinates and altitude (meters above sea level).

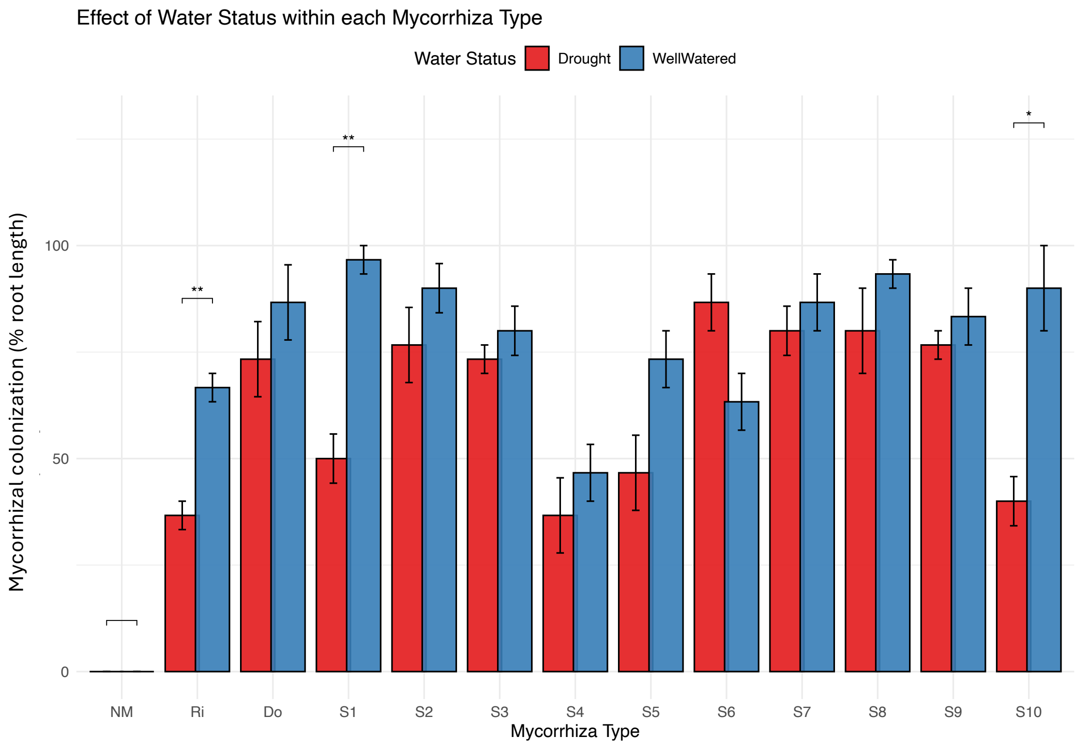


**Figure S2. Mycorrhizal colonization with native inocula under drought.** Mycorrhizal colonization was determined in plants grown under controlled conditions with native Moroccan inocula S1-S10 (see **Figure S1**). For comparison, inocula from *R. irregularis* DAOM197198 (Ri) and the drought adapated AM fungus *D. omaniana* (Do) were included, as well as non-mycorrhizal controls (NM). Plants were cultured for 8 weeks after inoculation under well-watered conditions (blue), or drought (red columns). Mean values (n=5) ± SE are shown. Significant effects ($P \leq 0.05$ one-way ANOVA and Tukey's test) of drought on root colonization are indicated * $P < 0.05$; ** $P < 0.01$; *** $P < 0.001$.

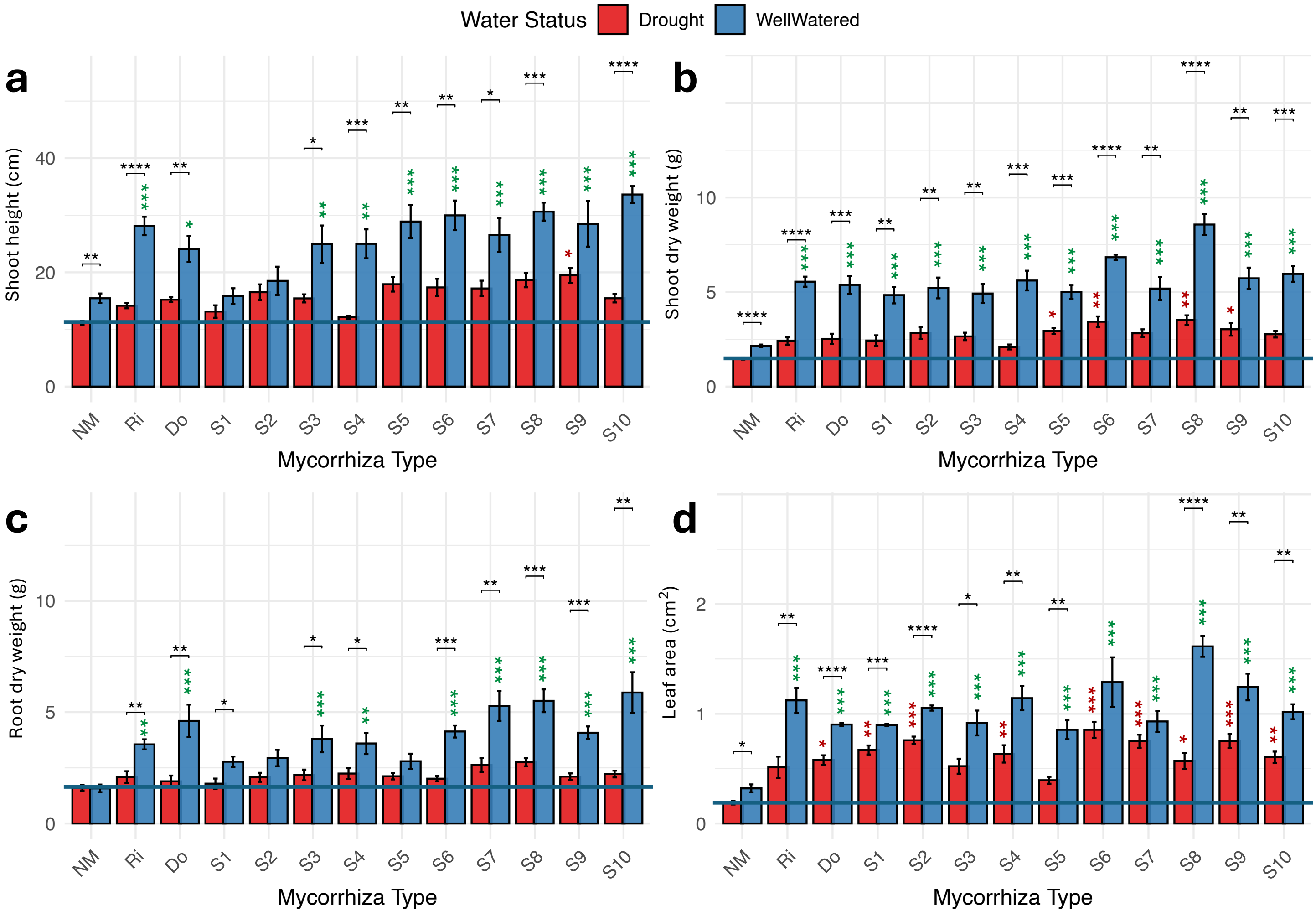


**Figure S3. Global growth parameters of plants colonized by native Moroccan AMF inocula.** Shoot height **(a)**, shoot dry weight **(b)**, root dry weight **(c)** and leaf area **(d)** of plants inoculated with 10 Moroccan inocula (S1-S10), and with the reference AMF species *R. irregularis* (RI) and *D. omaniana* (DO) and grown under well-watered or drought conditions, respectively. Horizontal lines indicate the level of non-mycorrhizal plants (NM) under drought as reference for the protective effect of the various mycorrhizal inocula under drought. Mean values (n=5) ± SE are shown. Significant effects of drought ($P \leq 0.05$; one-way ANOVA and Tukey's test) on the respective traits are indicated with black asterisks (* $P < 0.05$; ** $P < 0.01$; *** $P < 0.001$). Colored asterisks indicate significant differences of the inoculated plants vs. non-mycorrhizal controls under well-watered conditions (green asterisks), and under drought (red asterisks), respectively (* $P < 0.05$; ** $P < 0.01$; *** $P < 0.001$).

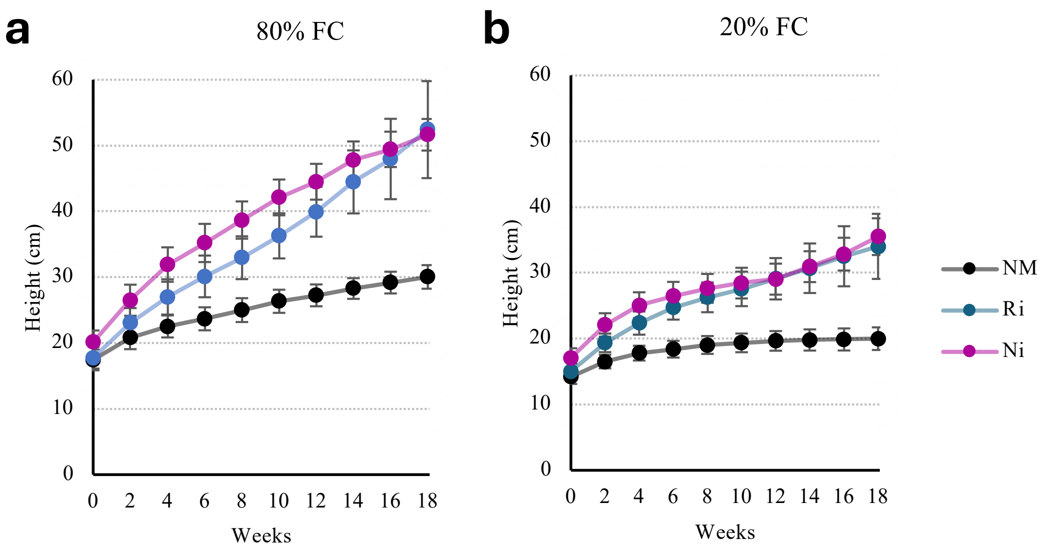


**Figure S4. Growth dynamics of argan colonized by *R. irregularis* under drought.**

Shoot height of plants inoculated with *R. irregularis* (Ri), with native inoculum S10 (Ni), or non-mycorrhizal (NM) under well-watered or drought conditions, respectively. Mean values (n=5) ± SE are shown.

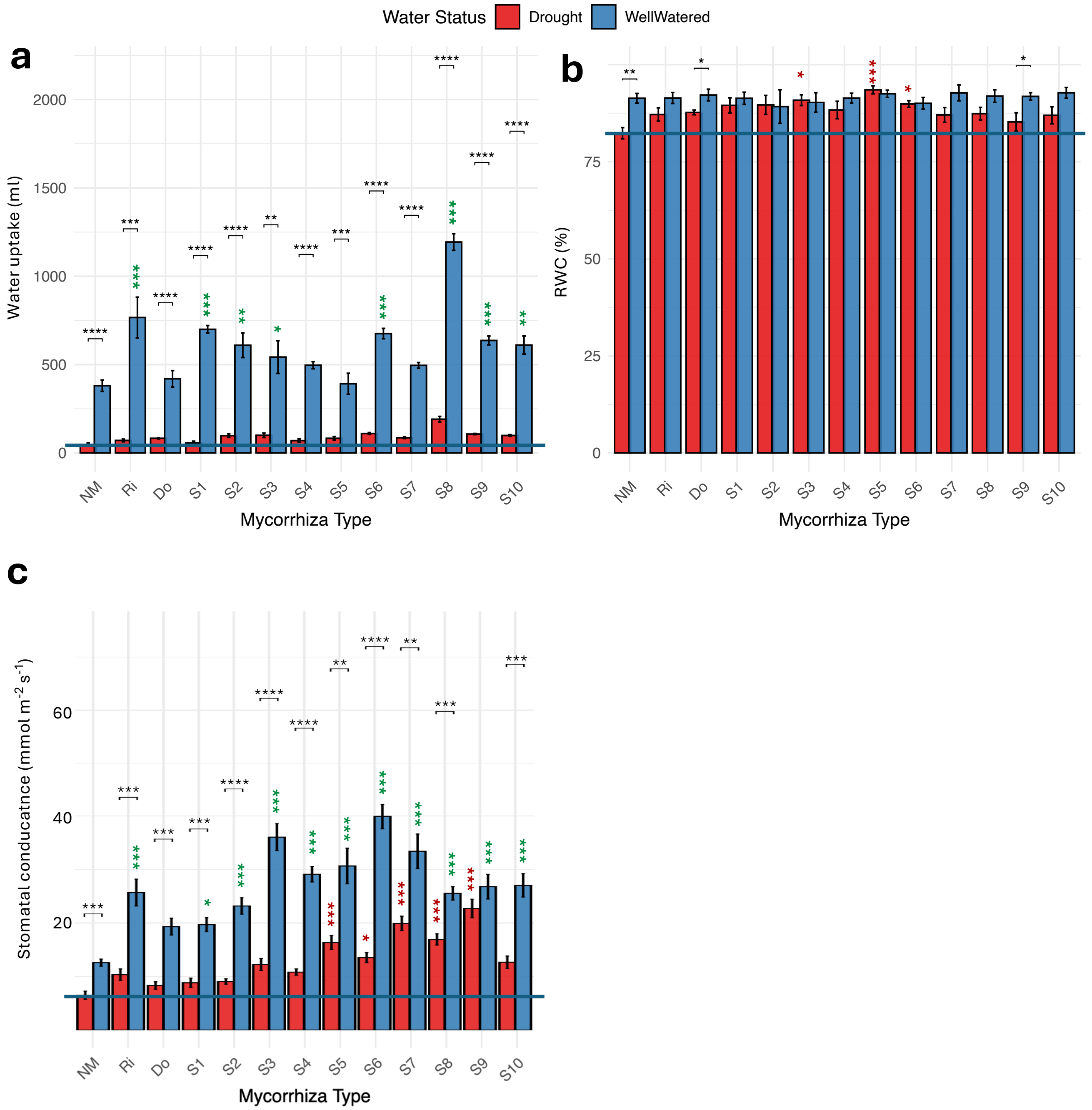


**Figure S5. Effects of mycorrhizal colonization on water relations.**
Accumulated water uptake **(a)**, leaf relative water content (RWC) **(b)**, and stomatal conductance **(c)** of plants inoculated with 10 Moroccan inocula (S1-S10), and with the reference AMF species *R. irregularis* (RI) and *D. omaniana* (DO) and grown under well-watered (blue columns) or drought conditions (red columns). Horizontal lines indicate the level of non-mycorrhizal plants (NM) under drought as reference for the protective effect of the various mycorrhizal inocula under drought. Mean values (n=5) ± SE are shown. Significant effects of drought ($P \leq 0.05$; one-way ANOVA and Tukey's test) on the respective traits are indicated with black asterisks (* $P < 0.05$; ** $P < 0.01$; *** $P < 0.001$). Colored asterisks indicate significant differences of the inoculated plants vs. non-mycorrhizal controls under well-watered conditions (green asterisks), and under drought (red asterisks), respectively (* $P < 0.05$; ** $P < 0.01$; *** $P < 0.001$).

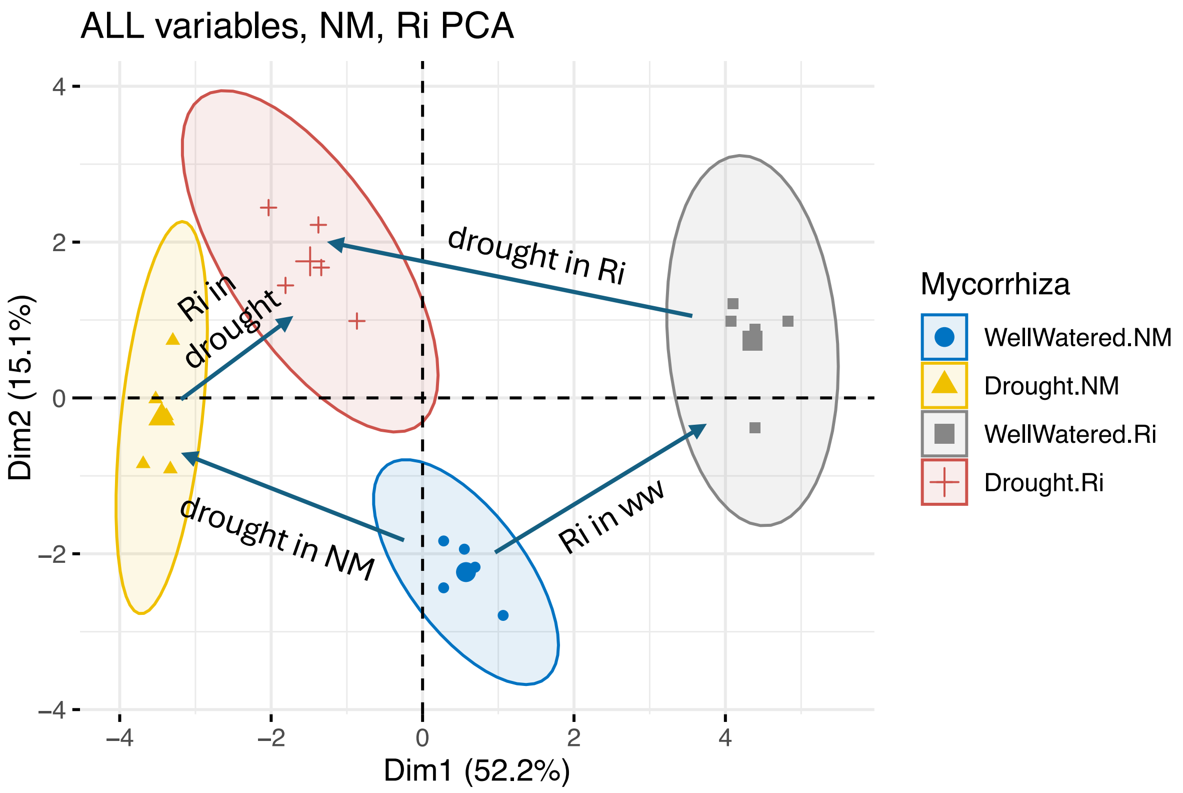


**Figure S6. Principle component analysis of phenotypic traits of plants with or without *R. irregularis* and under well-watered or drought conditions.**

Notable most effects were observed in the horizontal dimension (Dim1), which determined 52% of the effects. Mycorrhizal status caused a shift to the upper right, while drought caused an opposite shift to the left. Annotated arrows indicate in which direction the cloud as shifted by the respective factor.

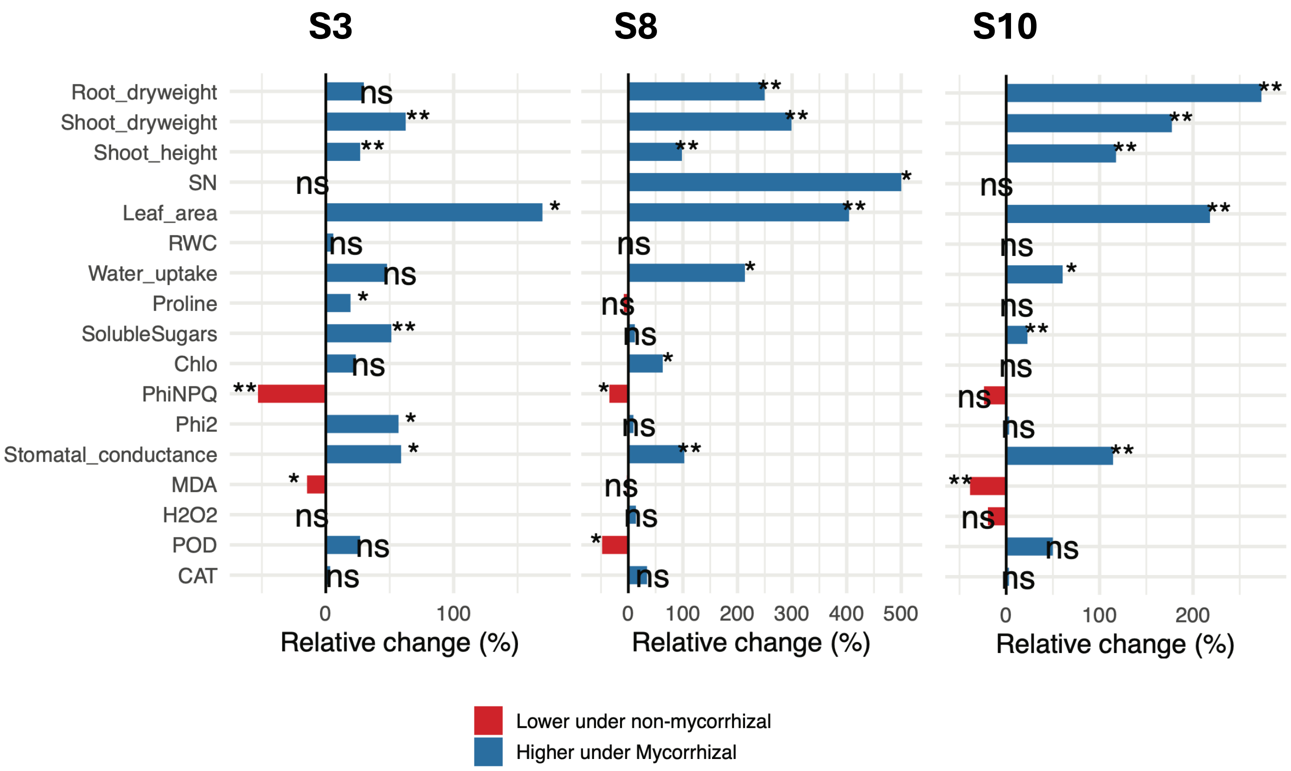


**Figure S7. Mycorrhizal effects on various growth-related parameters in well-watered plants inoculated with three native Moroccan inocula (S3, S8, S10).**
Relative difference (%) in 17 growth-related parameters determined in mycorrhizal plants vs. non-mycorrhizal controls grown under well-watered conditions. Eight weeks after inoculation with native inocula S3, S8 or S10, the following parameters were determined: root dry weight, shoot dry weight, shoot height, number of lateral branches (SN), leaf area, relative water content of the leaves (RWC), accumulated water uptake during the experiment, the content of proline, soluble sugars, and chlorophyll (Chlo) in the leaves, non-photochemical quenching (PhiNPQ), quantum yield of photosystem II (Phi2), the levels of malondialdehyde (MDA) and hydrogen peroxide (H2O2, as well as the activities of (POD), and catalase (CAT). Bars represent mean values (n=5); significance levels (one-way ANOVA and Tukey's test) are indicated as follows * $P < 0.05$; ** $P < 0.01$.

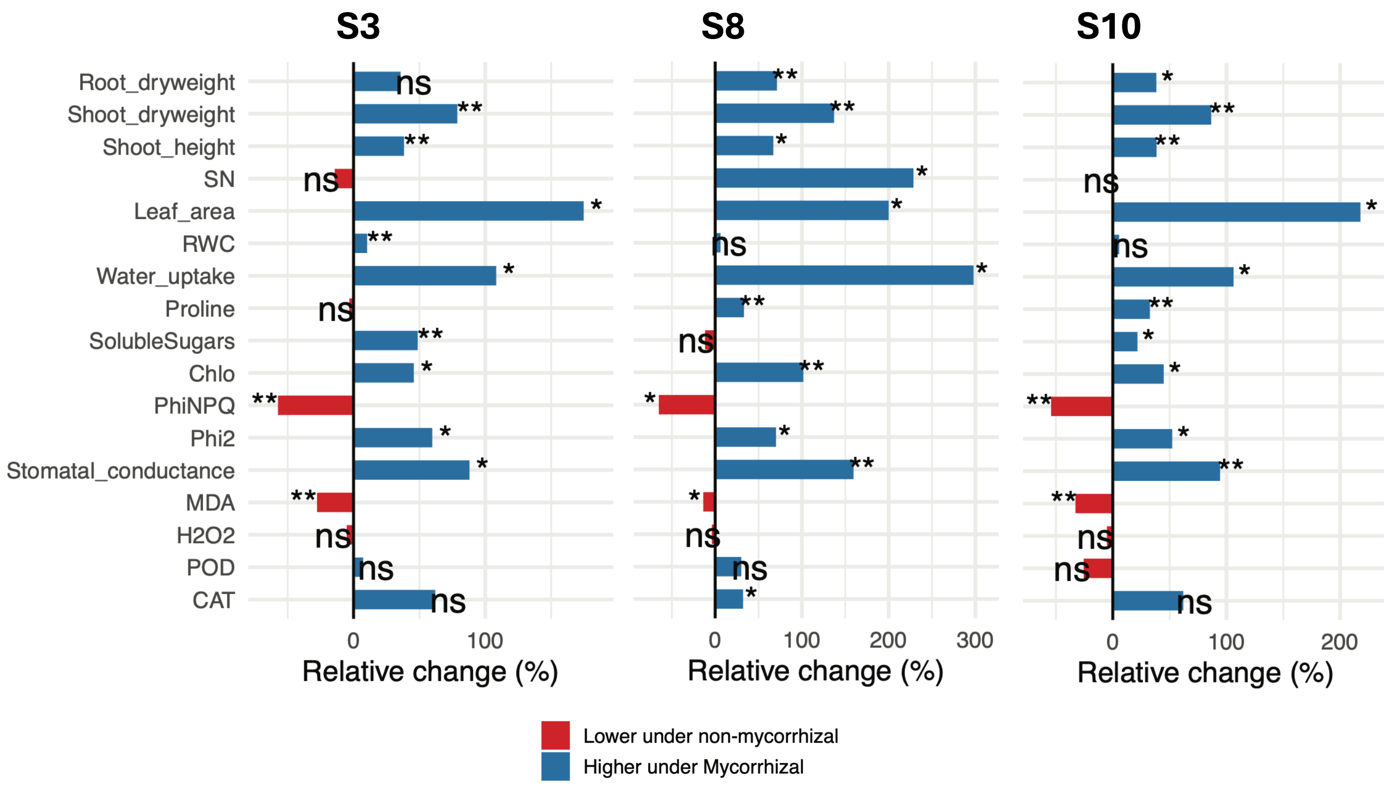


**Figure S8. Mycorrhizal effects on various growth-related parameters of plants inoculated with three native Moroccan inocula (S3, S8, S10) and grown under drought conditions.**

Relative difference (%) in 17 growth-related parameters determined in mycorrhizal plants vs. non-mycorrhizal controls grown under drought. Eight weeks after inoculation with native inocula S3, S8 or S10, the following parameters were determined: root dry weight, shoot dry weight, shoot height, number of lateral branches (SN), leaf area, relative water content of the leaves (RWC), accumulated water uptake during the experiment, the content of proline, soluble sugars, and chlorophyll (Chlo) in the leaves, non-photochemical quenching (PhiNPQ), quantum yield of photosystem II (Phi2), the levels of malondialdehyde (MDA) and hydrogen peroxide ($H_2O_2$, as well as the activities of (POD), and catalase (CAT). Bars represent mean values (n=5); significance levels (one-way ANOVA and Tukey's test) are indicated as follows * $P < 0.05$; ** $P < 0.01$.

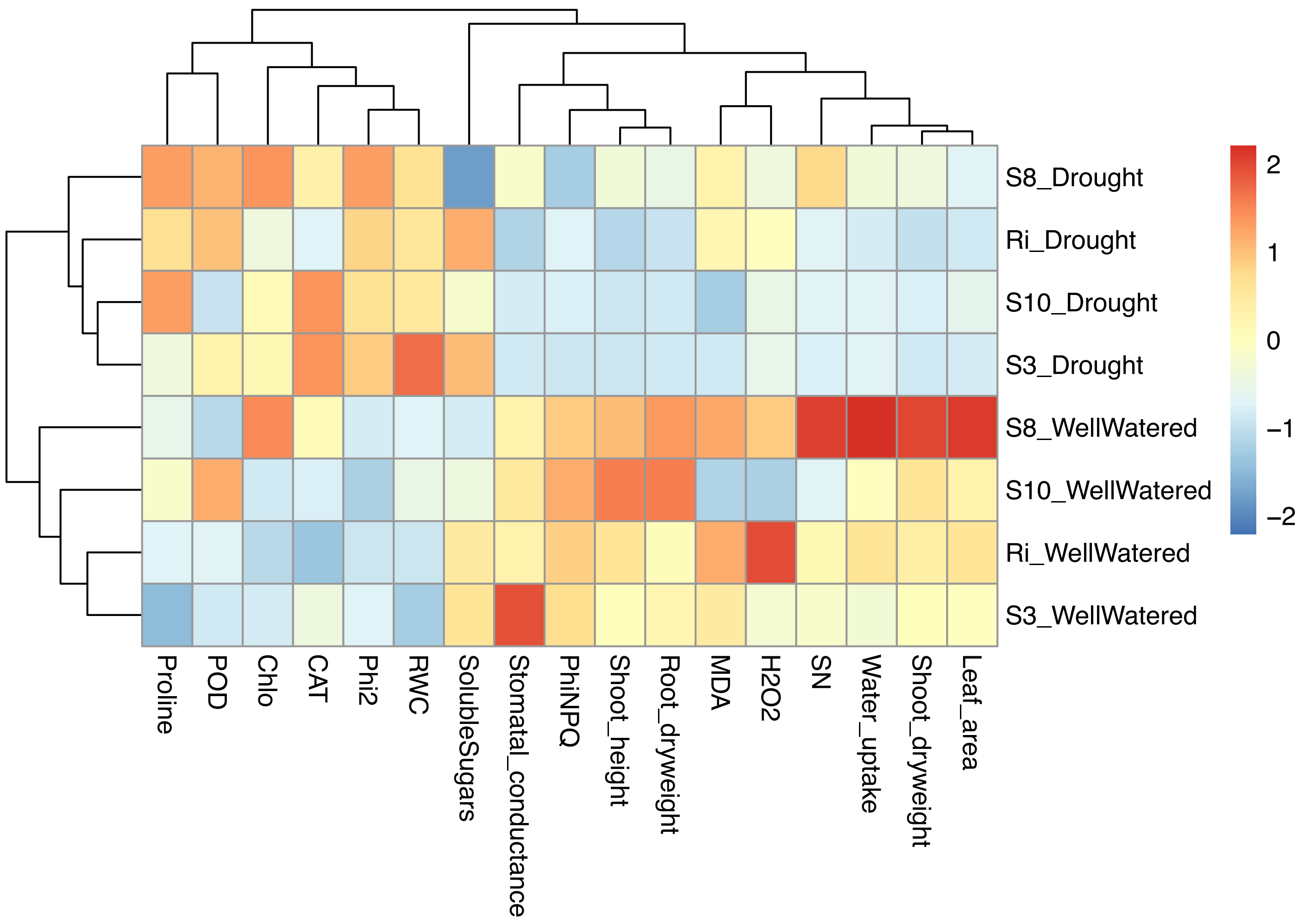


**Figure S9. Global clustering analysis of plants grown under the two water regimes with four different inocula.**

Heatmap shows fold change (log2) of the 17 growth- and stress-related parameters relative to the respective non-mycorrhizal controls. Note clustering according to water conditions, and the general reduction by drought of parameters related to growth, and a general induction of parameters related to stress.

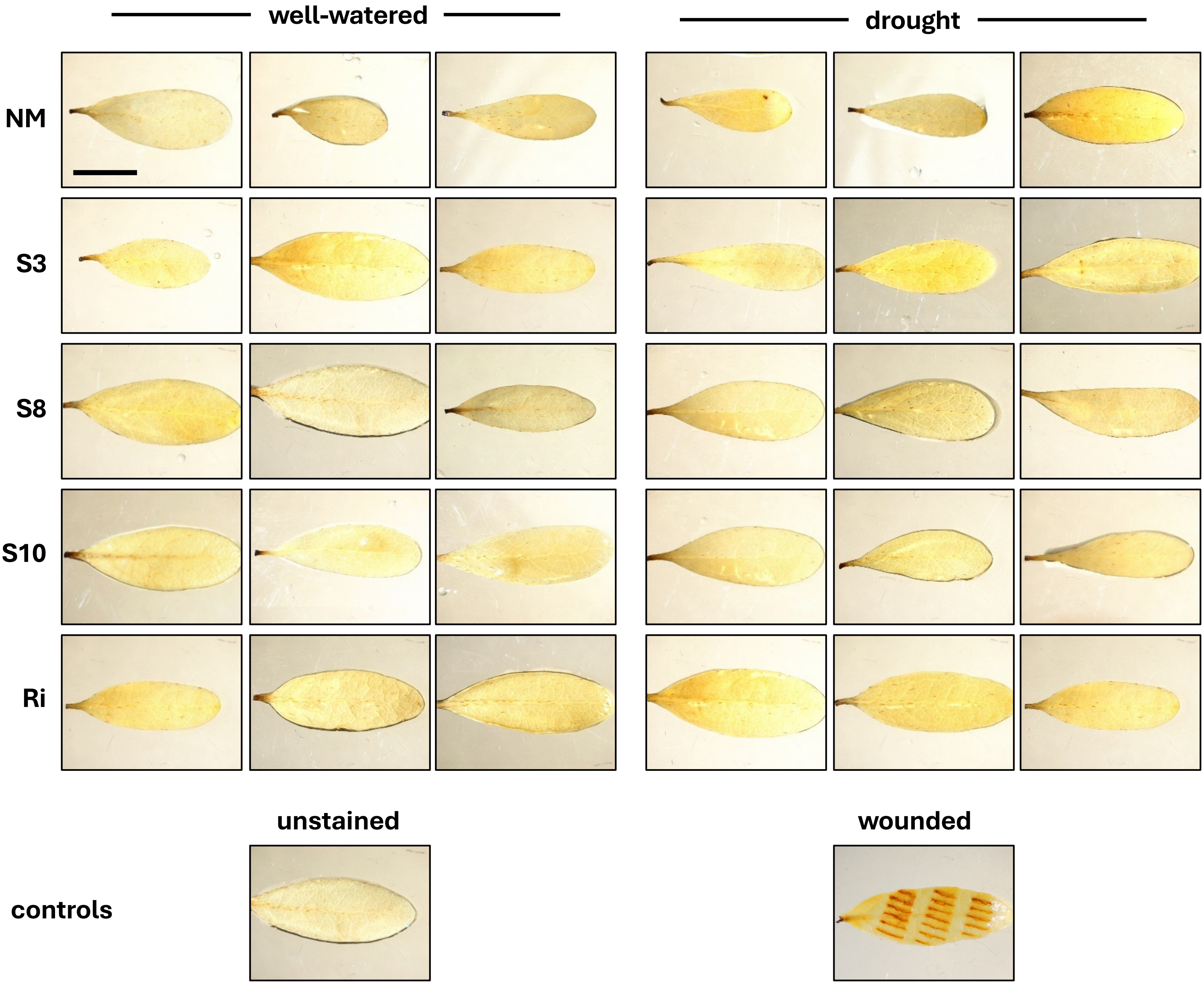


**Figure S10. Histochemical DAB staining of $H_2O_2$ in the leaves.**
Staining of leaves with diaminobenzidine (DAB) for $H_2O_2$ detection in argan plants grown under well-watered conditions and under drought, growing without AMF inoculum (NM), with three native inocula (S3, S8 and S10), or with *R. irregularis* (RI). No DAB staining was observed except in the positive control wounded by forceps.

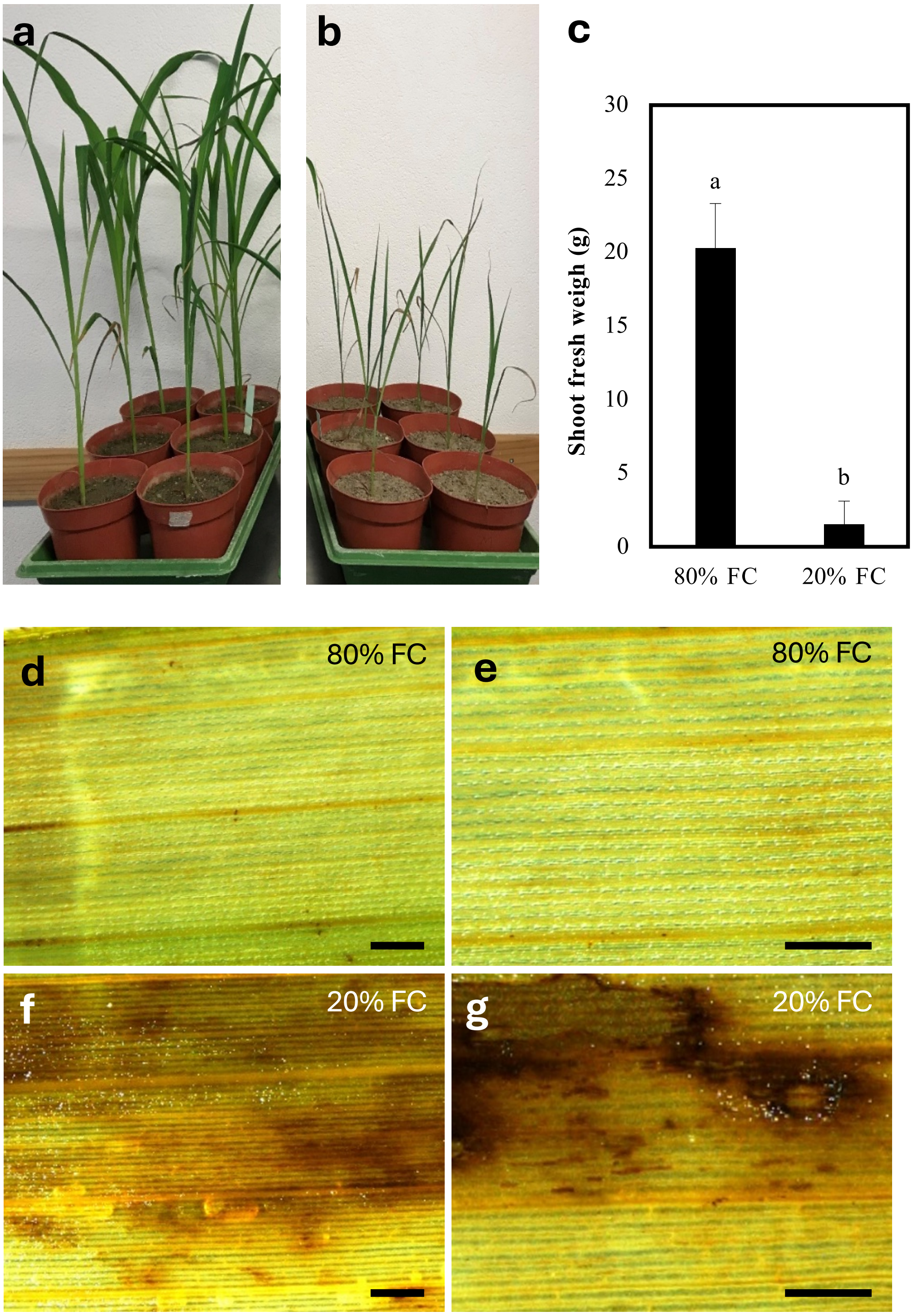


**Figure S11. Drought stress in maize.**

**(a-c)** Shoot appearance and fresh weight of maize plants grown under well-watered conditions **(a)**, and drought **(b)**, and respective shoot fresh weight **(c)**.

**(d-g)** Histochemical DAB staining of $H_2O_2$ in the leaves of maize plants grown under well-watered conditions **(d,e)** or drought **(f,g)**.

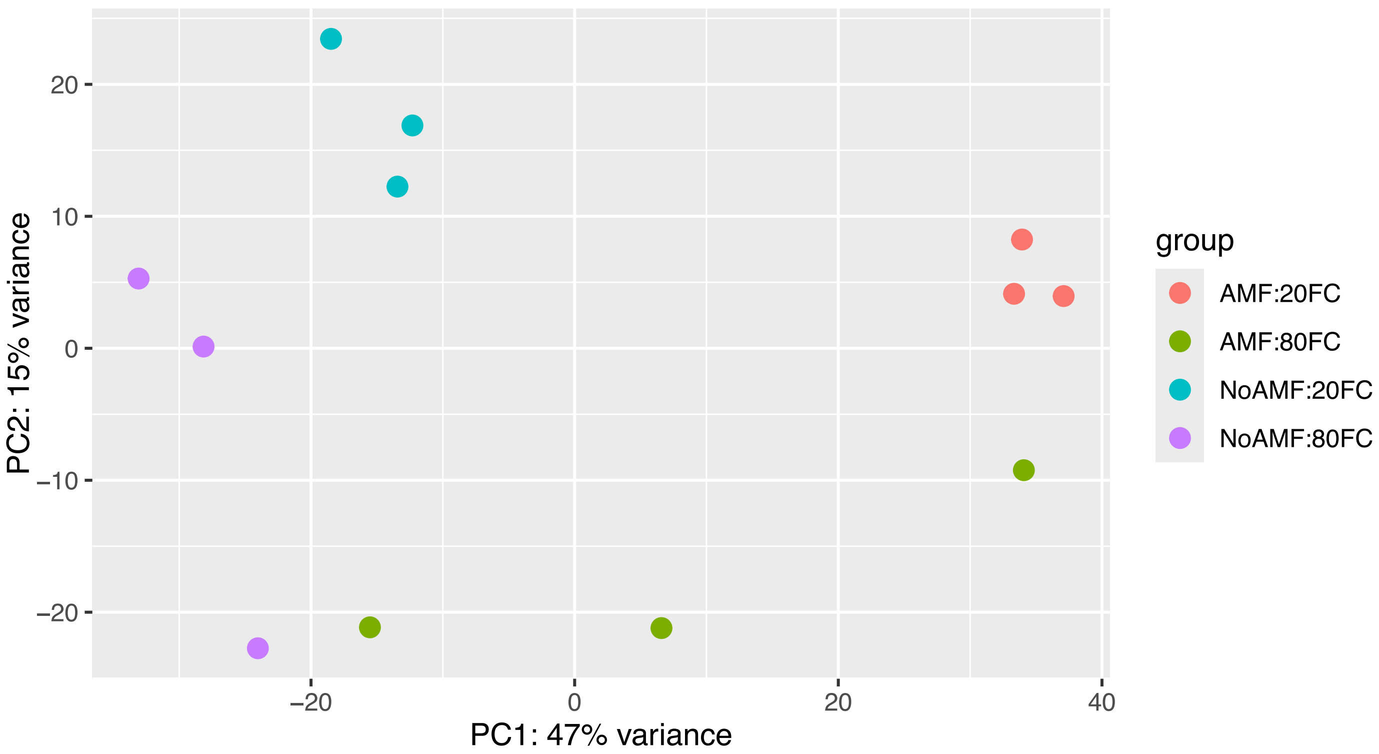


**Figure S12. Principle-Component analysis of RNAseq data.**
Plants wer grown with or without mycorrhizal inoculum (AMF, noAMF, respectively), under well-watered conditions (80% FC), or under drought (20% FC). Mycorrhizal samples appeared well clustered, while non-mycorrhizal samples were rather spread.

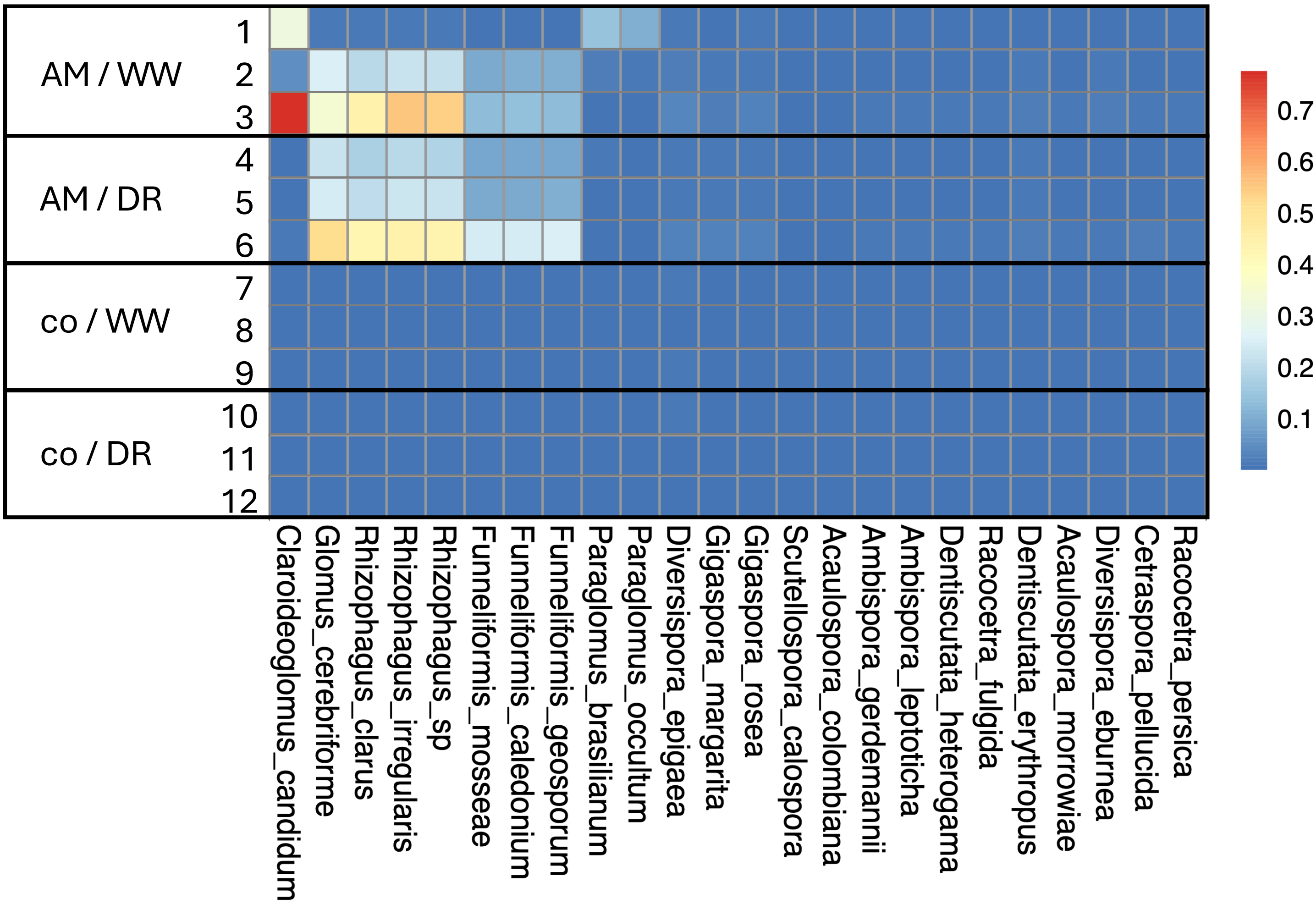


**Figure S13. AM fungal composition of inoculum S8.**
Fungal RNA-Seq reads were assigned to 24 AM fungal species for which a genome sequence was available. The heatmap indicates relative frequency. Fungal reads were restricted to inoculated plants (AM). Inoculum S8 contained mainly reads assigned to the genera Rhizophagus, Funnelliformis, Glomus, Claroideoglomus and Paraglomus.

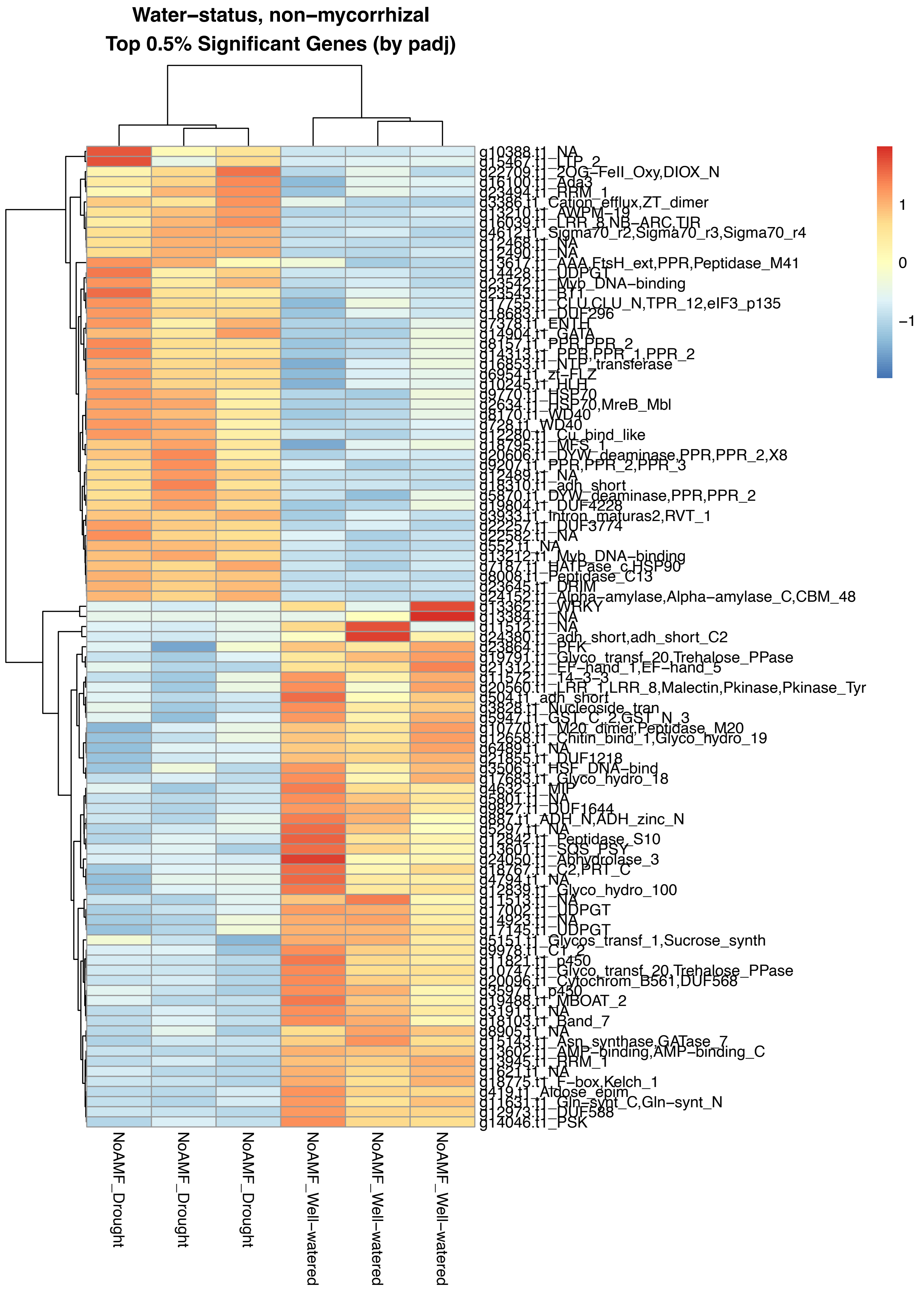


**Figure S14. Regulation of drought-responsive genes in non-mycorrhizal controls.** Clustering of the top 0.5% genes based on the p-values for significance of induction. Shown are genes induced or repressed by drought in non-mycorrhizal samples. The heatmap shows relative expression level.

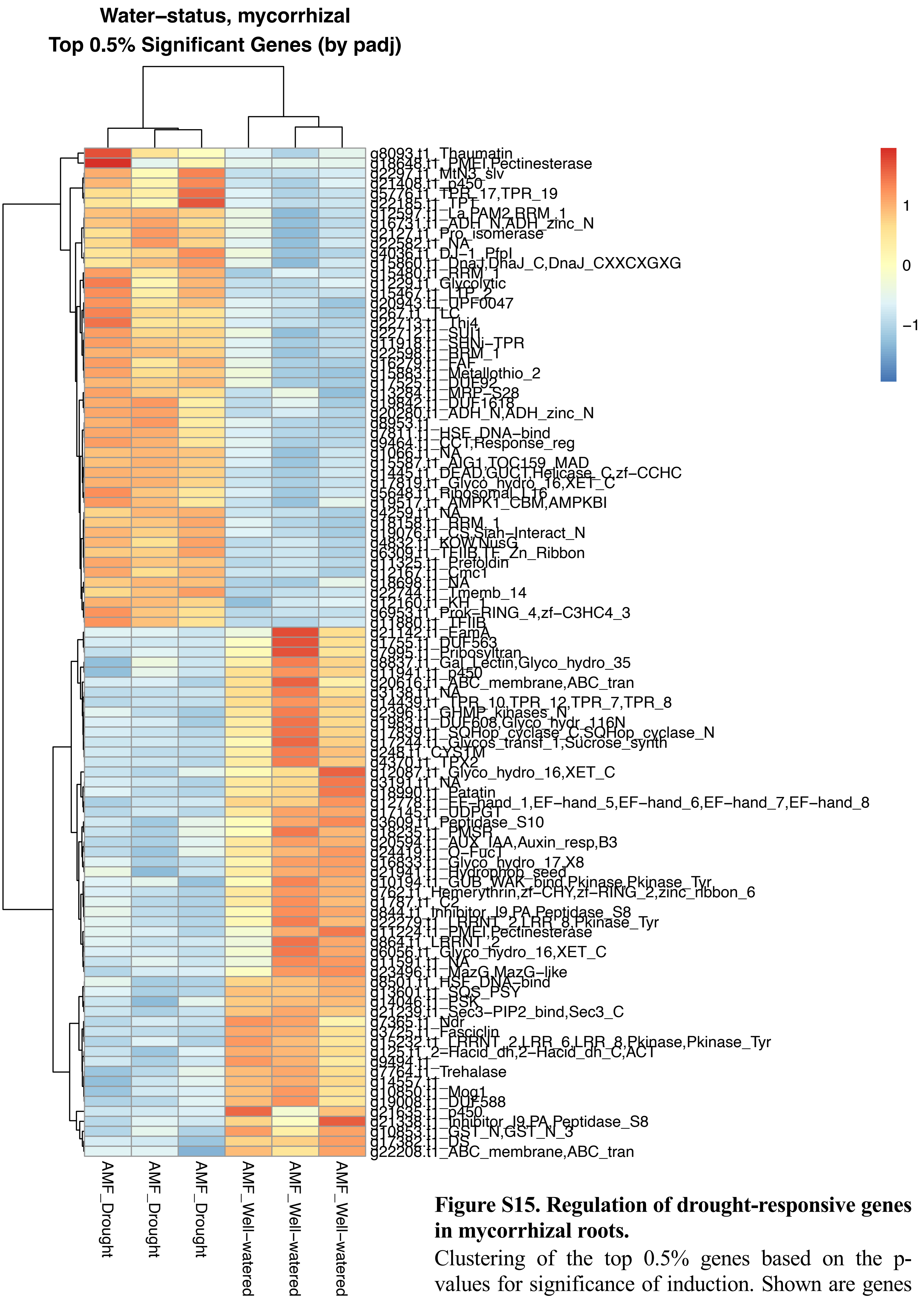


**Figure S15. Regulation of drought-responsive genes in mycorrhizal roots.**

Clustering of the top 0.5% genes based on the p-values for significance of induction. Shown are genes induced or repressed by drought in mycorrhizal samples. The heatmap shows relative expression level.

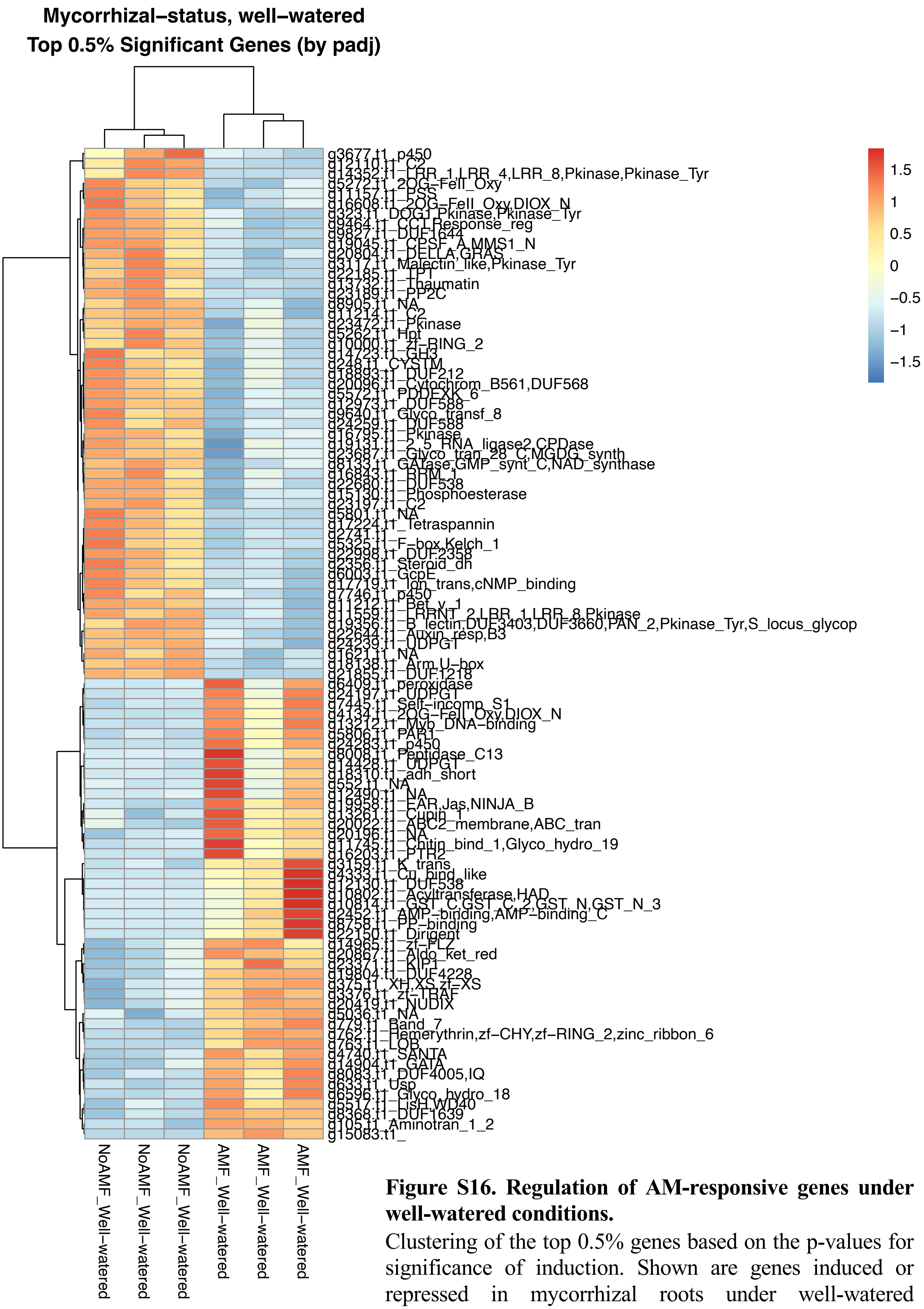


**Figure S16. Regulation of AM-responsive genes under well-watered conditions.**

Clustering of the top 0.5% genes based on the p-values for significance of induction. Shown are genes induced or repressed in mycorrhizal roots under well-watered conditions. The heatmap shows relative expression level.

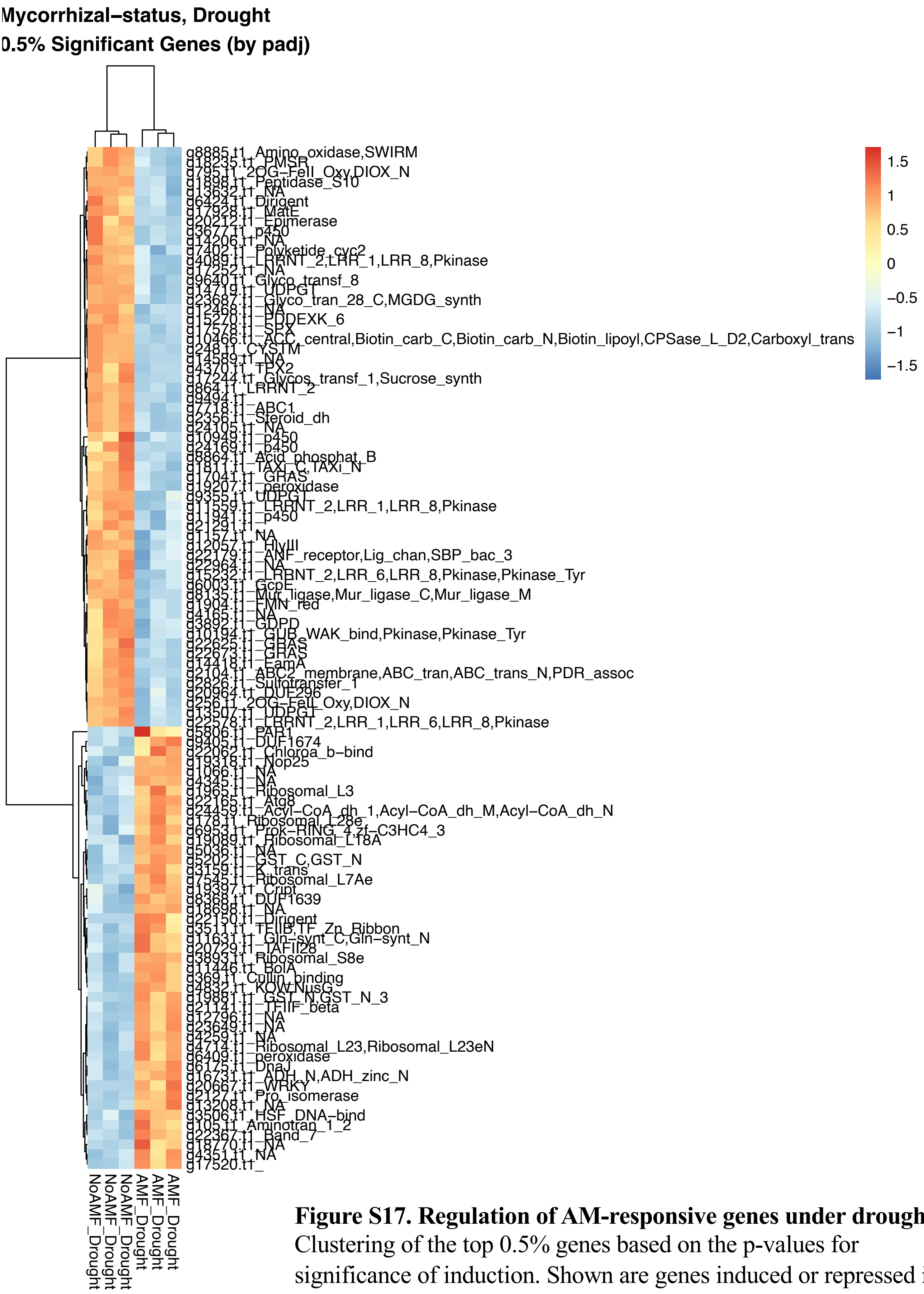


**Figure S17. Regulation of AM-responsive genes under drought.** Clustering of the top 0.5% genes based on the p-values for significance of induction. Shown are genes induced or repressed in mycorrhizal roots under drought. The heatmap shows relative expression level.

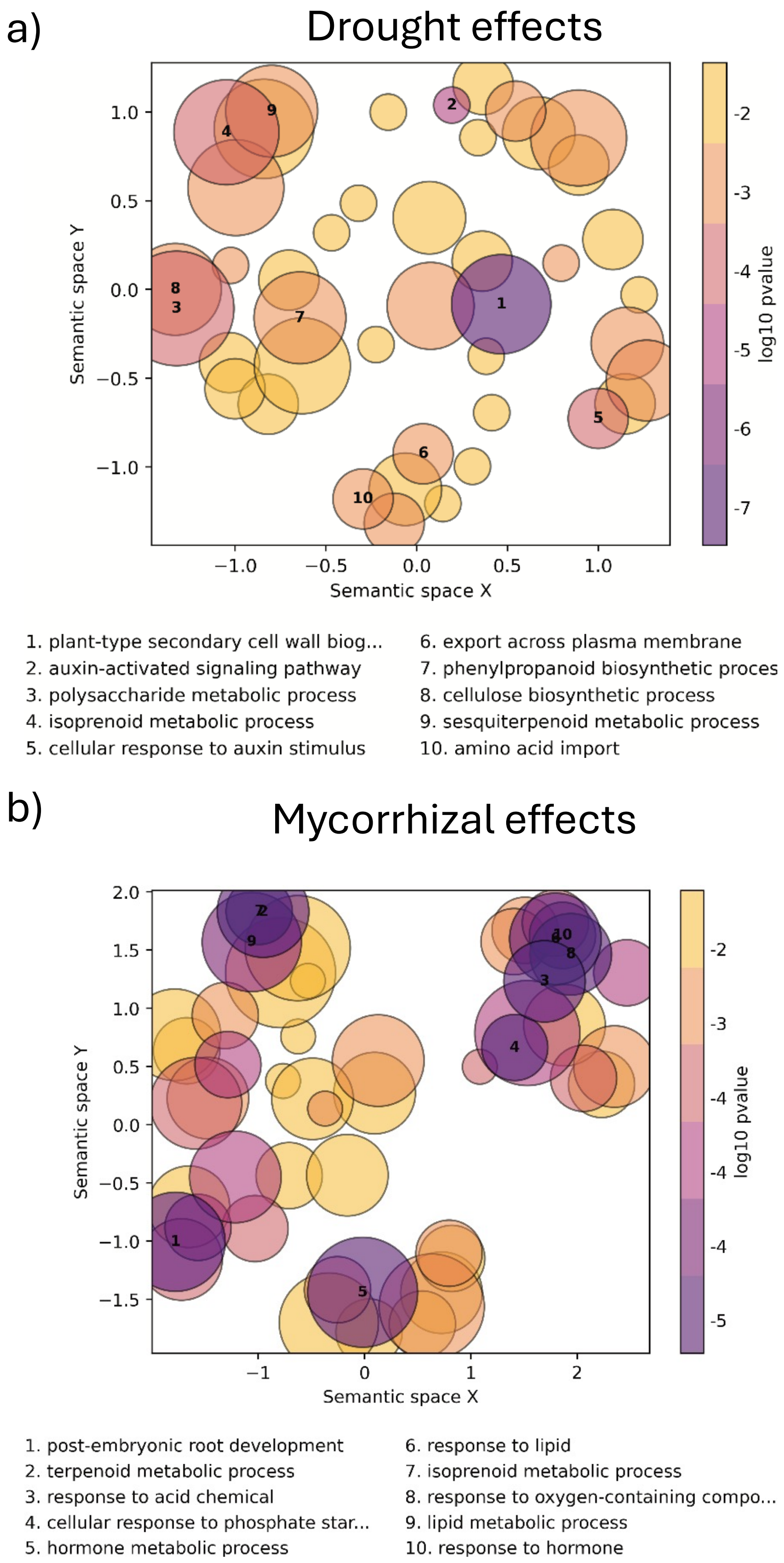


**Figure S18. Semantic space visualization of Gene Ontology (GO) term enrichment.** Scatter plots summarizing the enriched GO biological processes associated with the drought effect **(a)** and the mycorrhizal effect **(b)**, generated with GO-Figure. Enriched GO terms are projected into a two-dimensional semantic space (Semantic dimensions X and Y) based on the semantic similarity of their GO categories. Spatially adjacent points represent biological processes that are functionally closely related. The color of each bubble corresponds to the statistical significance of the enrichment. The size of the bubbles represent the size of each enriched GO-term.
